# Asymmetric centromere and gene locus positioning in *Drosophila* neural stem cells

**DOI:** 10.1101/2025.09.14.676095

**Authors:** Jennifer A. Taylor, Clemens Cabernard

**Affiliations:** Department of Biology, University of Washington, Seattle, WA 98105, USA

## Abstract

Chromatin organization is important for cell division, epigenetic memory, and gene regulation. It is often reflected in the non-random positioning of centromeres, but the underlying mechanisms and functions remain largely unknown. Here, we demonstrate that asymmetrically dividing *Drosophila* neural stem cells cluster centromeres near the nuclear envelope, adjacent to the apical centrosome. This asymmetric centromere positioning is regulated through microtubules, originating from the apical centrosome that connect to apical nuclear pore complexes. The minus-end directed motor protein Dynein, its binding partner Mushroom body defect, and Lamin are also required. Asymmetric centromere positioning persists throughout interphase in neural stem cells but is lost in more differentiated progeny. We also reveal that the genes *hunchback* (*hb*) and *pendulin* (*pen*) occupy specific nuclear regions, correlating with polarized centromere localization. We propose that fly neural stem cells translate their inherent polarity into stereotypical chromatin organization, potentially influencing cell fate decisions and stem cell behavior.

## Introduction

The spatial arrangement of chromatin in the interphase nucleus is highly organized and important for transcriptional regulation, DNA repair, and development ^1–4^. One manifestation of chromatin architecture is the position of centromeres, specialized regions on chromosomes that are enriched in repetitive DNA sequences. Centromeres serve as kinetochore protein assembly sites in mitosis, but their role in interphase is less understood ^5^. Centromeres can be positioned non-randomly within the nucleus. For instance, in 1885, it was first observed that centromeres cluster at one side of the interphase nucleus ^6,7^. In this arrangement, called Rabl configuration, centromeres are confined to one nuclear pole, opposite the nucleolus, and telomeres are dispersed at the opposite pole. The Rabl configuration has been found in yeast, plants, and vertebrate cells ^8–13^. In addition to the Rabl configuration, other non-random centromere positioning configurations have been described ^8^. For example, in *Drosophila* S2 cells, centromeres were found to be clustered around the nucleolus ^14^. In *Arabidopsis thaliana*, centromeres can form clustered foci at chromocenters, condensed regions of heterochromatin^15^. The function of non-random centromere positioning is mostly unclear, but it could limit the chromatin’s topological entanglement, support accuracy in gametogenesis, and aid in chromosome orientation and chromosomal integrity ^16–22^.

Non-random centromere organization can be implemented by linking centromeres to the cytoskeleton, thereby tying the spatial organization of the nucleus to larger-scale cellular organization. Several protein complexes have been implicated in connecting the cytoskeleton with chromatin. For instance, the Linker of Nucleoskeleton and Cytoskeleton (LINC) complex and the nuclear pore complex (NPC) have been shown to tether chromatin to the cytoskeleton ^23–26^. The LINC complex and NPCs have been implicated in centromere positioning. In *Schizosaccharomyces pombe*, the LINC complex protein Sad1 is required for Microtubule (MT)-dependent localization of the centromeres next to the Spindle Pole Body (SPB) ^27^. In *Arabidopsis*, both the LINC complex and the NPC are required for proper positioning of centromeres ^28,29^. However, the molecular mechanisms of non-random centromere positioning in metazoan cells remain to be defined.

Recently, we found that in larval *Drosophila melanogaster* neural stem cells (called neuroblasts, NBs), centromeres are non-randomly positioned throughout interphase ^30^. NBs are highly polarized, reflected in a molecularly distinct apical and basal cell cortex during mitosis. Dividing NBs retain the apical proteins while basal cell fate determinants segregate into either a differentiating Ganglion mother cell (GMC), or an Intermediate Progenitor Cell (INP) ^31,32^. Asymmetrically dividing NBs generate all the differentiated cells of the developing brain ^31,33^.

Neuroblasts also display clear polarization during interphase. For instance, neuroblasts maintain only one active microtubule-organizing center (MTOC) that is anchored to the apical neuroblast cortex ^34–36^. This apical MTOC is necessary to cluster centromeres close to the nuclear envelope, facing the active MTOC. Thus, biased MTOC activity is necessary for asymmetric centromere positioning ^30^. However, the molecular mechanisms connecting asymmetric centromere localization to cell polarity and how this form of chromatin organization is functionally important are unknown.

Here, we use asymmetrically dividing *Drosophila* NBs to define the mechanisms and function of centromere positioning in the developing larval brain. Using live cell imaging in the intact larval central nervous system, combined with detailed quantitative image analysis, we show that non-random and polarized centromere positioning is retained in self-renewing neuroblasts and their early, but not late, progeny. We further show that, unlike in *S. pombe* and *A. thaliana*, the LINC complex are dispensable for asymmetric centromere positioning. In contrast, we find that in NBs, the nucleoporins (Nups) Nup54, Nup62, and Nup98-96 are required for asymmetric centromere positioning. We also demonstrate that cytoplasmic heavy chain Dynein, the Nuclear Mitotic Apparatus (NuMA)-like protein Mushroom body defect (Mud), and Lamin are also required for asymmetric neuroblast centromere positioning. Finally, we show that the developmentally regulated gene loci *hunchback* (*hb*) and *pendulin* (*pen*) occupy specific areas within the nucleus relative to the apical polarity axis and that this organization requires polarized centromere localization. We propose a model whereby polarized centromere positioning is established and maintained by connecting MTs from the active apical MTOC to apically facing NPCs, using Dynein and Mud as a force-generating machinery. On the intranuclear side, NPCs either connect to centromeres directly or via Lamin. Collectively, this study suggests that polarized fly neural stem cells translate their intrinsic polarity into stereotypic chromatin organization, possibly regulating cell fate decisions and stem cell behavior.

## Results

### Neural stem cell centromeres form polarized clusters close to the apically facing side of the nuclear envelope

We previously reported that in neuroblasts (NBs) of third instar *Drosophila* larval brains, centromeres retain a polarized localization near the apical centrosome throughout interphase ^30^. To gain mechanistic insight into this polarized centromere positioning, we performed time-lapse imaging and quantitative image analysis in intact third instar larval brains, expressing the centromere-specific Histone H3 variant Centromere Identifier (Cid; CENPA in humans) tagged with EGFP (EGFP::Cid) ^37^, as well as the microtubule marker UAS-mCherry::α-Tubulin ^38^. While expression of UAS-mCherry::α-Tubulin is induced via the neuroblast-specific worGal4 driver ^39^, EGFP::Cid expression is regulated via its endogenous regulatory elements. Since α-Tubulin is excluded from the nucleus during interphase, we were able to use the α-Tubulin signal to segment both the apical centrosome (AC) and the nucleus using a machine learning algorithm (Fig. 1a; Movie 1; see Methods for details). For the quantitative analysis, we selected NBs that progressed through a complete interphase and defined the start of interphase as the time point when the nuclear envelope had reformed after cytokinesis. The end of interphase was defined as the time point when the α-Tubulin signal in the nucleus had increased as a result of nuclear envelope breakdown (NEB). To pool measurements from multiple cells and to account for different lengths of interphase, we binned measurements based on the percent of interphase elapsed (see Methods for details).

**Figure 1:**
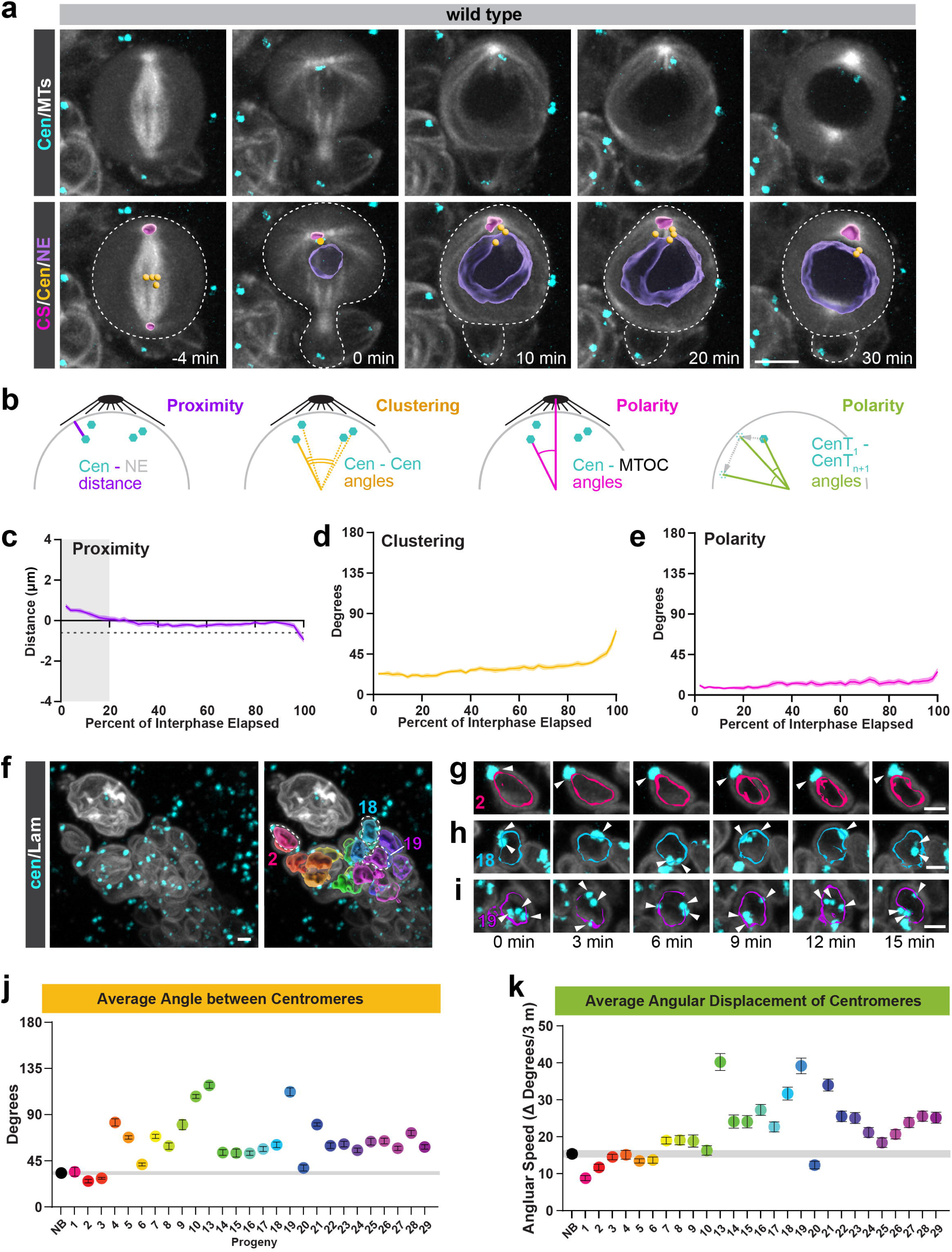
Centromeres are polarized, clustered, and peripherally localized in interphase NBs. **(a)** Top row: representative third instar larval NB expressing worGal4, UAS-mCherry::α-Tubulin (white) and centromere marker Cid::EGFP (cyan) from metaphase through prophase. 0 minutes is defined as the beginning of nuclear envelope reformation. Bottom row: computational segmentations of the centrosome (CS; magenta), nucleus (NE; purple), and centromeres (Cen; yellow). Dotted lines delineate cell outline of the NB and newborn progeny cell. **(b)** NB segmentation was used to quantify (1) proximity, measuring the centromere distance to the nearest point on the nuclear envelope (Cen – NE distance); (2) clustering, measuring the angle between individual centromere pairs (Cen – Cen angles); (3) polarity, measuring the angle between centromeres and the apical microtubule organizing center (MTOC; Cen – MTOC angles). For differentiating progeny, lacking a defined MTOC, polarity was quantified by measuring the angle between centromeres for all (n) time points of the movie (Cen T1 – CenT1+n). **(c)** Centromere – nuclear envelope proximity plot. Negative value indicates that the centromere is positioned inside the segmented nucleus while a positive value indicates that the centromere is outside. The dotted line indicates the 15% radial distance boundary. Values outside of this region are no longer considered to be in close proximity to the NE. Inaccuracies in nuclear periphery segmentation, which occur in late cytokinesis/early interphase, are highlighted with a grey bar. **(d)** Centromere clustering and **(e)** centromere polarity plots. All measurements are from a total of 18 NBs from 3 brain lobes. Average values at each timepoint are represented by the bold line. Shaded region indicates the 95% confidence interval. **(f-i)** Representative NB clone expressing worGal4, UAS-Lamin::GFP (white) and the centromere marker Cid::mRFP (cyan). Computational reconstructions (right image) of progeny nuclei are shown in rainbow colors according to assigned progeny order. **(g-i)** Image sequences of three example progeny cells. White arrowheads indicate the positions of centromeres. Numbers indicate assigned birth order. **(j)** The average angle between pairs of centromeres over the course of the movie for the NB and progeny #1-29, excluding dividing cells #11 and 12. Bars indicate the 95% confidence interval. The shaded gray box indicates the 95% confidence interval for the NB. **(k)** The average angular speed of centromeres (measured as the magnitude of the change in degrees per 3 minutes) measured over the course of the movie for the NB and progeny #1-29, excluding #11 and 12. Bars indicate the 95% confidence interval. The shaded gray box indicates the 95% confidence interval for the NB. Scale bars: 5 µm in (a), 2 µm in (f,i). Time in minutes (min).

As previously reported ^30^, we confirmed that centromeres appear to be positioned at the nuclear periphery (Fig. 1a). To quantify this, we measured the distance between centromeres and the closest point on the surface of the segmented nucleus throughout interphase (Fig. 1b). Throughout interphase, centromeres in wild-type NBs were positioned on average -0.08 µm (SD= +/-0.41 µm) away from the nuclear envelope (Fig.1c; Fig. S1a). Considering that the average third instar neuroblast radius is 4.05 µm (SD = +/-0.74 µm), this puts centromeres within 2% proximity of the nuclear surface (Fig. S1d). To account for inaccuracies in NE segmentations and the observed variations in NB nuclear size, we defined centromeres that were positioned within 15% proximity (≥ 0.61 µm for NBs) of the nuclear surface as peripherally localized. Based on this definition, wild-type centromeres are peripherally localized during the majority of interphase (96%) but start moving away from the periphery before NEB (Fig. 1c; Fig. S1a; Movie1).

We also noticed that in wild-type NBs, centromeres form clusters, with Cid-positive spots grouped together. To quantify this clustering, we calculated the average angle among centromeres throughout interphase (Fig. 1b and methods). We found that for the majority of interphase (68%), the average angle between centromeres remained below 30°. At the end of interphase (74% of interphase elapsed and later), the average angle started to increase above 30° (Fig. 1d; Fig. S1b; Movie1).

We previously reported that interphase centromeres appear to be positioned near the apical centrosome (AC) ^30^. To quantify this polarization, we determined the centromere’s position and calculated the average angle between the centromeres and the AC throughout interphase (Fig. 1b; Fig. S1c). For the majority of interphase (98%), the average angle between the apical AC and centromeres remained below 20°. As expected from the decrease in centromere clustering before NEB, we detected a subtle increase in the average angle at the end of interphase (Fig. 1e; Fig. S1c; Movie1). Thus, centromeres stay polarized toward the AC throughout interphase. In conclusion, these measurements show that in wild-type interphase NBs, centromere positioning is (1) peripheral, (2) clustered, and (3) polar.

### As NB progeny mature, centromeres lose clustering and polarity

To determine if the peripheral, clustered, and polarized centromere arrangement is NB-specific, we analyzed centromere positioning in NB progeny. NBs divide asymmetrically, generating a self-renewed NB and either a small Ganglion mother cell (GMC) or Intermediate Progenitor (INP) with every division. INPs mature and divide asymmetrically a few times, generating GMCs. GMCs formed from NBs or INPs undergo a terminal symmetric division, producing two neurons or glia ^31^. Thus, central brain neuroblast lineages allow a comparison of centromere positioning between stem and differentiating cells. More importantly, since all cells within a NB lineage are derived from the parental NB, the position of INPs, GMCs, and neurons reflects their birth order: cells generated more recently are positioned closer and more mature cells further away from the parental NB. To quantify centromere positioning in differentiating cells, we labelled individual NB lineages with UAS-lamin::EGFP ^40^ (driven by worGal4), co-expressing Cid::mRFP ^41^, and segmented all nuclei and centromere spots (see Methods for details). We assigned approximate lineage order to the cells by multiplying the distance between the progeny and NB centroids times the angle between the progeny centroid and NB division axis (Fig. 1f – i). For simplicity, we also excluded dividing progeny (#11 and #12) from our measurements. We measured centromere – nuclear envelope proximity and clustering in NBs and progeny as before (Fig. 1b). We found that in both young (cell # 1-10) and older offspring (cell # 13-29), centromeres are localized on average 0.18 µm (young: SD= +/-0.27 µm; older: SD= +/-0.28 µm) from the segmented nuclear surface (Fig. S1f). Considering that the nuclear radius of differentiating cells is on average 1.49 µm (SD= +/-0.17 µm; Fig. S1e), centromeres are within 15% of the nuclear envelope, indicating that centromeres are also peripherally localized in differentiating progeny cells.

Early offspring cells (cells 1, 2, 3) had clustering angles below 35° (cell 1: mean = 34.46°, SD= +/-34.16°. Cell 2: mean = 25.36°, SD= +/-15.07°. Cell 3: mean = 28.15°, SD= +/-8.75°), similar to the clustering angle in the NB (mean = 33.05°; SD= +/-12.62°). However, the clustering angle in nearly all later offspring (cells # 4-29) was significantly increased (Fig. 1j; Fig. S1g).

We observed that centromere positioning in relation to the first time point varied considerably among differentiating progeny. While centromeres remained relatively stationary in young progeny, we observed a lot more centromere movement in cells further away from the parental NB (Fig. 1g-i). Since most progeny cells do not contain a discernible MTOC (see below and ^34^), we used the change in centromere angular positioning over time to determine if centromeres are polarized (Fig. 1b). Centromere angular movement in progeny #1-6 is less than that of the NB, but most of the progeny #7-29 show significantly increased centromere movement (Fig. 1k). These measurements show that early offspring retain peripheral, clustered, and polarized centromere positioning, while clustering and polarization is lost in older progeny.

To determine if the change in centromere positioning correlates with a loss of the interphase MTOC, we stained third instar larval brains containing clones expressing UAS-mCherry::αTubulin under the control of worGal4 with the pericentriolar matrix (PCM) marker anti-centrosomin (Cnn), the nuclear envelope marker anti-Lamin, and the centromere marker anti-Cid. Since PCM is necessary to nucleate microtubules and to form an active MTOC ^42^, we analyzed all cells in the marked lineage for Cnn localization in non-mitotic cells. We observed that in the vast majority of interphase progeny, no MTOCs were detectable. The two exceptions we observed were in the progeny cell immediately adjacent to the NB (Fig. S1h) and sometimes in a pair of cells not immediately adjacent to the parental NB (Fig S1i). In the latter case, the MTOCs were positioned at opposing ends of each cell, suggesting that they are remnants of a recent mitosis. Older progeny further away from the parental Nb showed no discernable PCM signal (Fig.S1j). Taken together, we conclude that differentiating progeny lose centromere clustering and polarization, most likely due to the lack of functional MTOCs.

### Centromere positioning requires an active MTOC

As we previously reported, polar centromere positioning in NBs is dependent on microtubules from the apical MTOC ^30^. Interphase NBs contain a single, active, apically localized MTOC ^34–36,43–52^. The centriolar protein Centrobin is a key protein in maintaining apical MTOC activity ^43,48,49^. To precisely define the role of MTOCs in centromere positioning, we imaged live third instar larval NBs expressing Cnb-RNAi and mCherry::α-Tubulin controlled by worGal4, together with endogenously expressing Cid::EGFP. In *cnb* RNAi NBs, the apical centrosome disappeared in mid-interphase (defined as when the AC can no longer be segmented) and reappeared in prophase (Fig. 2a; time points 15 min and 43 min, respectively; Movie2). Centromeres in *cnb* RNAi showed a slight but statistically significant decrease in peripheral positioning but remained within 15% of the radial distance to the nuclear periphery until the end of interphase (Fig. 2b; Fig. S4a). Clustering was unaltered compared to wild type during the majority of interphase (Fig. 2c; Fig. S4a). However, consistent with recently published results ^30^, loss of Cnb caused a significant increase in the angle between the AC and the centromeres following loss of the AC (Fig. 2d; Fig. S4a).

**Figure 2:**
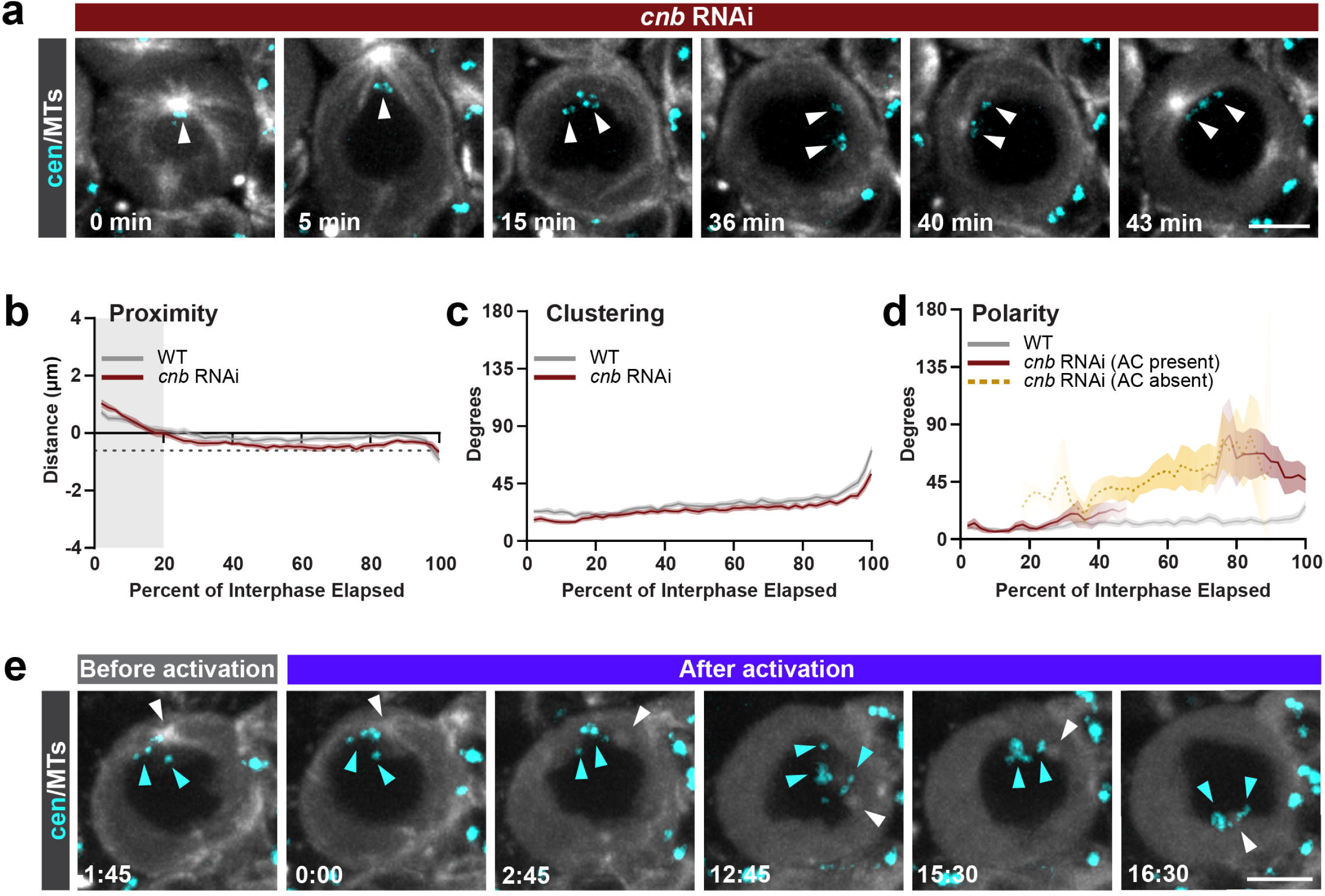
Centromere positioning is microtubule-dependent. **(a)** Image series of representative *cnb* RNAi third instar larval NB expressing worGal4, UAS-mCherry::α-Tubulin (white) and centromere marker Cid::EGFP (cyan) from the beginning of nuclear envelope reformation (t=0) through prophase. White arrows indicate the positions of centromeres. Graphs of the **(b)** distance from the nuclear periphery to the centromeres (dotted line = 15% radial distance; gray bar represents inaccurate nuclear periphery segmentation), **(c)** angle between each pair of centromeres, and **(d)** angle between the apical centrosome and the centromeres for n = 13 cells from N=2 brain lobes. The lines indicate the average at each timepoint and the shaded areas represent the 95% confidence interval for *cnb* RNAi (dark red) and wild type (gray). After the apical centrosome disappeared during interphase in *cnb* RNAi, measurements between the last known position of the apical centrosome and the current position of the centromeres were calculated and plotted separately (gold, dotted line). **(e)** Wild-type third instar NB treated with the photoactivatable microtubule inhibitor SBTub2M ^53^, which was activated with a pulse of UV light (t=0). White arrows indicate the positions of centrosomes, and teal arrows indicate the position of centromeres. Scale bars = 5 µm. Time in minutes (min).

To further probe the contribution of microtubules to centromere positioning, we treated cells with the previously described photoactivatable microtubule inhibitor SBTub2M ^53^. We observed that after activation of the inhibitor with a pulse of UV, microtubules disappeared, and most MTOCs were reduced to a faintly discernible spot. While unperturbed apical MTOCs retain their position at the apical cell cortex throughout interphase ^43–45,49^, we observed that in a subset of NBs exposed to activated SBTub2M, MTOC remnants moved away from the apical side of the cell but remained in close proximity to the NE. In each of these cases, the centromere clusters tracked along with the MTOC remnant (Fig. 2e). These data suggest that while the microtubule-based connections of the MTOC to the apical cortex were lost, the MTOC remnant remained connected to the centromeres. Taken together, we conclude that the connection of microtubules from the apical MTOC to the apical cortex is necessary for positioning centromeres to the apically facing side of the nucleus throughout interphase.

### Nuclear pore complexes are required for NB centromere positioning

Next, we investigated the mechanism of NB centromere positioning. We hypothesized that microtubules from the apical MTOC must be linked to the nucleoplasmic centromeres. Linkages between microtubules and the nucleus are commonly facilitated by members of the LINC complex (LInker of Nucleoskeleton and Cytoskeleton), which contain SUN or KASH domains ^54,55^. Therefore, we assessed whether members of the *Drosophila* LINC complex are necessary for centromere polarity. NBs, deficient for the SUN-domain protein Klaroid (Koi; koi^80^ (null allele) / *koi* deficiency), as well as NBs expressing RNAi against *koi,* still positioned centromeres close to the apical MTOC (Fig. S2a, b). Similarly, NBs lacking the KASH-domain protein Klarsicht (Klar) retained polarized centromere distribution (Fig. S2c-f). And *msp300* mutant NBs, obtained from crossing a null allele of the KASH-domain protein Muscle-specific protein 300 kDa *(Msp300; msp300*^Δ*KASH*^ ^56^ ) to a *msp300* deficiency chromosome (Msp300; *msp300*^Δ*KASH*^ */ deficiency*), also retained their polarized distribution (Fig. S2g).

In male germline stem cells, centromeres are connected to microtubules crossing the NE in early prophase ^57^. To test whether microtubules from the apical NB MTOC also directly cross the NE during interphase, we performed transmission electron microscopy (TEM) of ultrathin sections of wild-type NBs. Microtubules were visible as two parallel lines spaced approximately 25 nm apart. The apical MTOC could be identified by microtubules radiating out from a central position, even if the centrioles were not visible in that section. While AC microtubules were often seen closely approaching the NE, we never observed microtubules crossing or inside the NE. However, we saw numerous examples of microtubules passing close to nuclear pore complexes (NPCs), which were visible as electron-dense barrel- or ring-like structures embedded in the NE. We observed microtubules oriented approximately parallel to the NPCs or approaching end-on (Fig. 3a-i).

**Figure 3:**
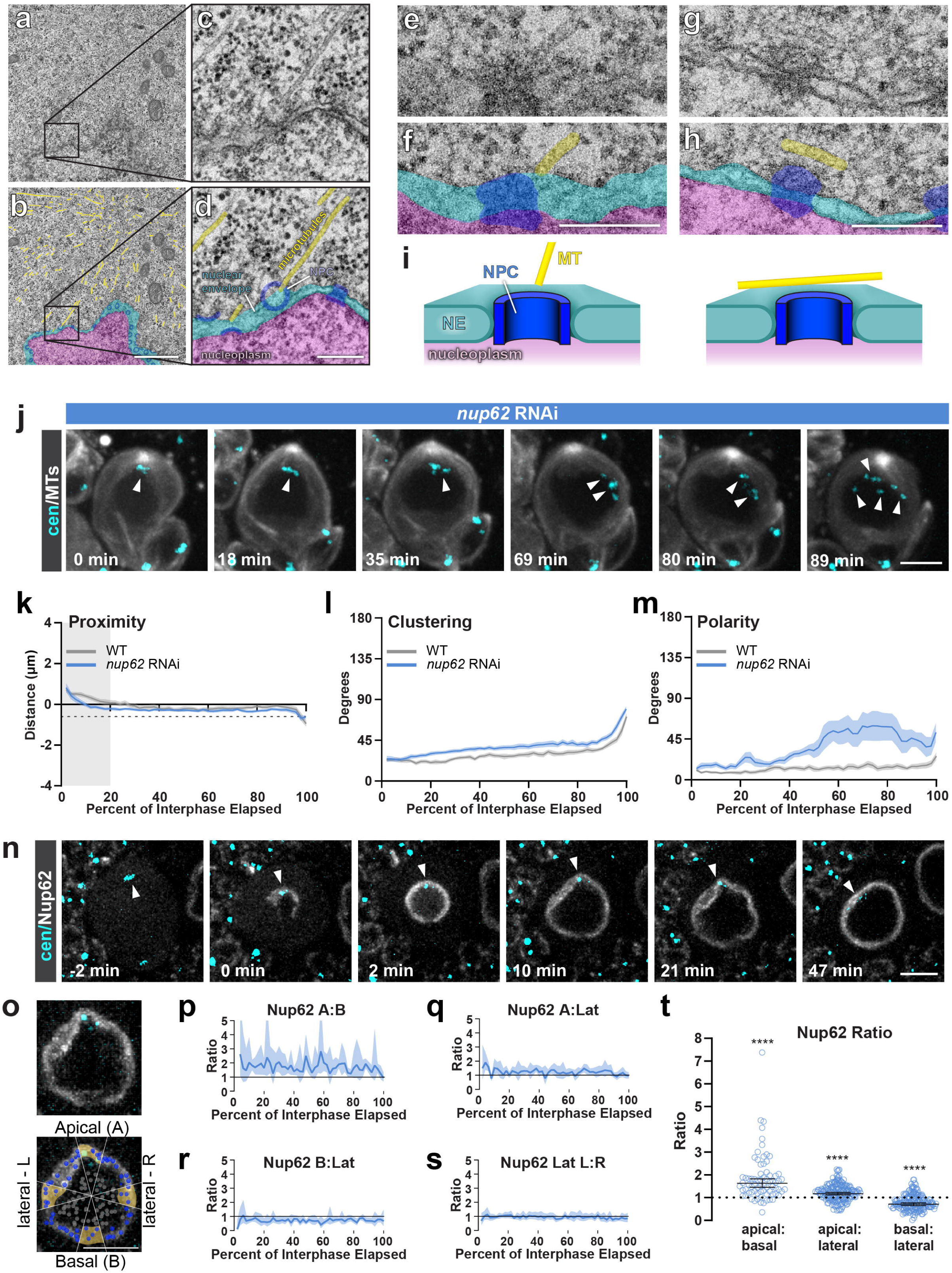
Nuclear pore complexes are required for NB centromere positioning. **(a-h)** Transmission electron micrographs of ultrathin sections of wild-type third instar larval NBs with magnified views (high magnification images in c, d correspond to the boxed regions in a,b) and colored overlay panels showing microtubules (yellow), the nuclear envelope (cyan), nuclear pore complexes (dark blue), and the nucleoplasm (magenta). **(e-h)** High-magnification images of apical centrosome microtubules in close proximity to nuclear pore complexes. **(i)** Schematic representation of observed microtubule orientations in relation to NPCs. **(j)** Representative image sequence of a *nup62* RNAi third expressing third instar larval NB co-expressing worGal4, UAS-mCherry::α-Tubulin (white) and the centromere marker Cid::EGFP (cyan) from the beginning of nuclear envelope reformation (t=0) through prophase. White arrows indicate the positions of centromeres. Graphs of the **(k)** distance from the nuclear periphery to the centromeres (dotted line = 15% radial distance; gray bar represents inaccurate nuclear periphery segmentation), **(l)** angle between each pair of centromeres, and **(m)** angle between the apical centrosome and the centromeres for n=13 cells from N=4 brain lobes. The lines indicate the average at each time point and the shaded areas represent the 95% confidence interval for *nup62* RNAi (blue) and wild type (gray). **(n)** Representative third instar larval neuroblast expressing EGFP::Nup62 (white) and the centromere marker Cid::mRFP (cyan) from telophase through the end of interphase, with t=0 being the beginning of nuclear envelope reformation. White arrows indicate the positions of centromeres. **(o)** Diagram of the spots used for Nup62::EGFP quantification in (p-t). Blue spots = nuclear surface spots used for Nup62::EGFP signal intensity measurements, gray spots = nucleoplasmic spots averaged and used for background subtraction at eact timepoint. Yellow regions indicate the angular regins used for apical, lateral, and basal signal quantification. **(p-s)** Graph of the ratio of (p) apical:basal, (q) apical:lateral, (r) basal:lateral, and (**s**) lateral left:right signal over the course of one interphase for the cell shown in (n). (t) Scatterplot of each ratio of Nup62::EGFP signal per timepoint over the course of one interphase for n = 5 cells. **** = p<0.0001; ratio t-test. Bar = geometric mean with 95% confidence interval in brackets. (a,b) Scale bars: (a, b): 1 µm; (c-h) 0.2 µm, (j, n): 5 µm, (q,r): 5 µm.

Since the NPC traverses the NE, we reasoned that it might function as a molecular bridge to link microtubules to centromeres. The NPC is a large, multimolecular complex composed mostly of a family of proteins known as Nucleoporins (Nups)^58^. To test this hypothesis, we assessed the contribution of Nups to centromere positioning by knocking down individual Nups using inducible RNAi (driven by *worGal4*). Knocking down Nup54, Nup62, and Nup96-98 caused a loss of centromere polarization without a loss of apical MTOC activity (Fig. 3j, Fig. S3a,b; Movie3). We proceeded with a more detailed investigation of Nup62 and confirmed its phenotype with an independent Nup62 RNAi line (data not shown) as well as in third instar *nup62^ex161^* homozygous null mutant NB clones (*nup62* null mutants do not survive to third instar stages). While wild-type control NB clones contained normal MTOCs and polarized centromeres, *nup62* mutant NB clones phenocopied the *nup62* RNAi phenotype, albeit with less penetrance (Fig. S3c, d).

To analyze Nup62’s phenotype in more detail, we measured centromere peripheralization, clustering, and polarity in *nup62* RNAi-expressing NBs. While loss of Nup62 mildly affected the peripheral localization of centromeres in early and late interphase, centromeres remained within 15% of the radial distance to the nuclear periphery (Fig. 3k, Fig. S4b). We also measured a small but significant decrease in centromere clustering, manifested as an increase in the inter-centromere angles for all but the beginning of interphase (8.2° change on average) (Fig. 3l, Fig. S4b). Most notably, we measured a large and significant increase in the angle between centromeres and the apical centrosome throughout interphase, with the largest difference in the second half of interphase (Fig. 3m, Fig. S4b).

Nup62 is a component of the NPC’s central channel ^59^, and we hypothesized that Nup62-containing NPCs could be enriched in the NE-portion close to centromeres, facing the apical MTOC. To answer this, we imaged NBs expressing Nup62::EGFP (donated to BL from Ch. Lehner). As previously reported for Nup58 and Nup107 ^60^, Nup62::EGFP was recruited first to the apical side of the nuclear envelope at mitotic exit (Fig. 3n; Movie4), but then proceeded to be incorporated throughout the nuclear envelope. We measured Nup62::EGFP intensity in the NE and quantified the ratio of the Nup62::EGFP signal on the apical vs. basal, apical vs. lateral, and basal vs. lateral sides of the nuclear envelope throughout interphase. These measurements showed a significant enrichment of Nup62::EGFP in the apical-facing NE region compared to lateral and basal NE regions (Fig. 3o-t).

Taken together, we conclude that Nup62, Nup54, and Nup98 are required for polarized centromere localization. Furthermore, the proximity of MTs to NPCs suggests that a connection between MTs and apical Nup62+ NPCs, rather than individual nucleoporins, regulates centromere localization.

### Lamin is required for both centromere polarity and clustering in NBs

Since Lamin has been shown to anchor NPC positions ^61–63^, we hypothesized that it might be required for apical centromere localization. Alternatively, it could be involved in centromere proximity and or clustering. To investigate this, we imaged *lamin* mutant NBs and measured centromere positioning. In *lam^A^*^25^*/lam^04643^* mutant NBs, centromeres showed small but statistically significant deviations from wild type peripheral localization. However, centromeres in *lamin* mutant NBs remained within 15% radial position of the nuclear periphery throughout interphase (Fig. 4a,b; Fig. S4c). Centromere clustering was significantly reduced for all but the beginning of interphase (18.4° change on average) (Fig. 4c; Fig. S4c). The angle between centromeres and the apical centrosome was similar to wild type at the very beginning of interphase but subsequently increased and remained significantly elevated for the rest of interphase (Fig. 4d; Fig. S4c; Movie5). We conclude that Lamin is necessary for both centromere clustering and polarization and has a subtle impact on peripheralization.

**Figure 4:**
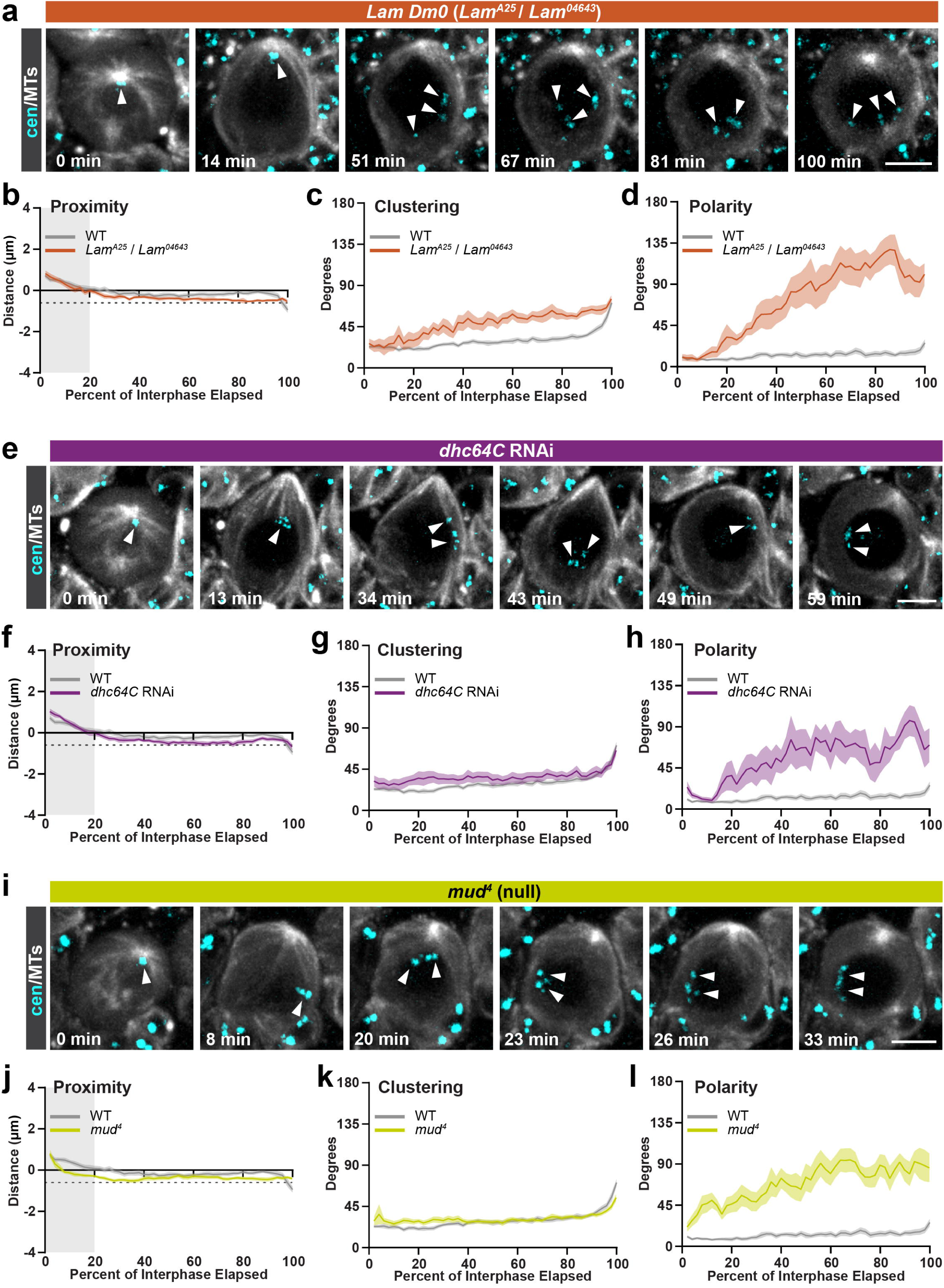
Lamin, cytoplasmic Dynein and the NuMA-like protein Mushroom body defect (Mud) are required for NB centromere positioning. Image series of representative **(a)** *Lam^A25^*/*Lam^04643^*, **(e)** *dhc64C* RNAi expressing, **(i)** *mud^4^* mutant third instar larval NB expressing worGal4, UAS-mCherry::α-Tubulin (white) and centromere marker EGFP::Cid (cyan) from the beginning of nuclear envelope reformation (t=0) through prophase. Graphs of the **(b, f, j)** distance from the nuclear periphery to the centromeres (dotted line = 15% radial distance; gray bar represents inaccurate nuclear periphery segmentation), **(c, g, k)** angle between each pair of centromeres, and **(d, h, l)** angle between the apical centrosome and the centromeres. The lines indicate the average at each timepoint, and the shaded areas represent the 95% confidence interval. n = 7 cells, N=2 brain lobes for *Lam^A25^*/*Lam^04643^;* n=8 cells, N=4 brain lobes for *dhc64C* RNAi; n = 12 cells, N=3 brain lobes for *mud^4^*. Scale bars: (a, e, i) bar = 5 µm,

### Dynein is required for centromere polarity in NBs

Microtubules can be linked to NPCs via molecular motors ^64^. We reasoned that a minus-end-directed motor protein might provide a pulling force at the NE to anchor centromeres to the AC. To test this hypothesis, we knocked down the heavy chain subunit of the cytoplasmic Dynein motor complex, Dynein heavy chain 64C (Dhc64C), with worGal4 inducible RNAi. We observed a loss of centromere polarity in cells that retained an apical MTOC throughout interphase (Fig. 4e; Movie6). We measured a significant increase in the average angle between the AC and centromeres for the majority of interphase (Fig. 4h; Fig. S4d). While centromeres displayed minor but statistically significant changes in peripheral localization compared to wild type, they remained within 15% radial distance of the nuclear periphery until the end of interphase (Fig. 4f; Fig. S4d). We also detected a subtle but significant increase in the average centromere clustering angle for part of interphase (6.7° change on average) (Fig. 4g; Fig. S4d). Thus, we conclude that cytoplasmic Dynein has a small impact on peripheral localization and clustering but is necessary for polarized centromere positioning.

### The NuMA-like polarity protein Mushroom body defect (Mud) is required for proper centromere positioning in NBs

The Nuclear Mitotic Apparatus (NuMA)-related protein Mushroom body defect (Mud) interacts with cytoplasmic Dynein and is necessary for maintaining the orientation of the mitotic spindle during mitosis ^65^. *mud* mutant neuroblasts contain normal apical-basal neuroblast polarity ^66–68^ and maintain an interphase MTOC ^49^. We investigated centromere localization in *mud*^4^ null mutants ^69,70^ and found that centromeres retained peripheral localization (within 15% radial position) and clustering (1.8° change on average). However, centromere polarization was compromised in *mud*^4^ mutant neuroblasts, with a significantly increased angle between centromeres and the AC throughout interphase (Fig. 4i-l; Fig. S4e; Movie7)

To investigate if Mud’s contribution to centromere localization is due to its interaction with Dynein, we measured centromere positioning in *mud* mutants lacking the Calponin homology (CH; *mud*^Δ*CH*^ ^71^) domain, which is located at the N-terminus (Fig. S5a). In NuMA, the CH domain has been shown to specifically fold as a Hook domain, binding directly to Dynein light intermediate chain, and is postulated to serve as an activator of cytoplasmic Dynein ^72^. Centromere positioning in *mud*^Δ*CH*^ phenocopied the *mud*^4^ phenotype; centromeres were positioned at the nuclear periphery (within 15% radial position until the end of interphase), remained clustered (4.8° change on average), but lacked polarization (Fig. 5a-d; Fig. S5b).

**Figure 5:**
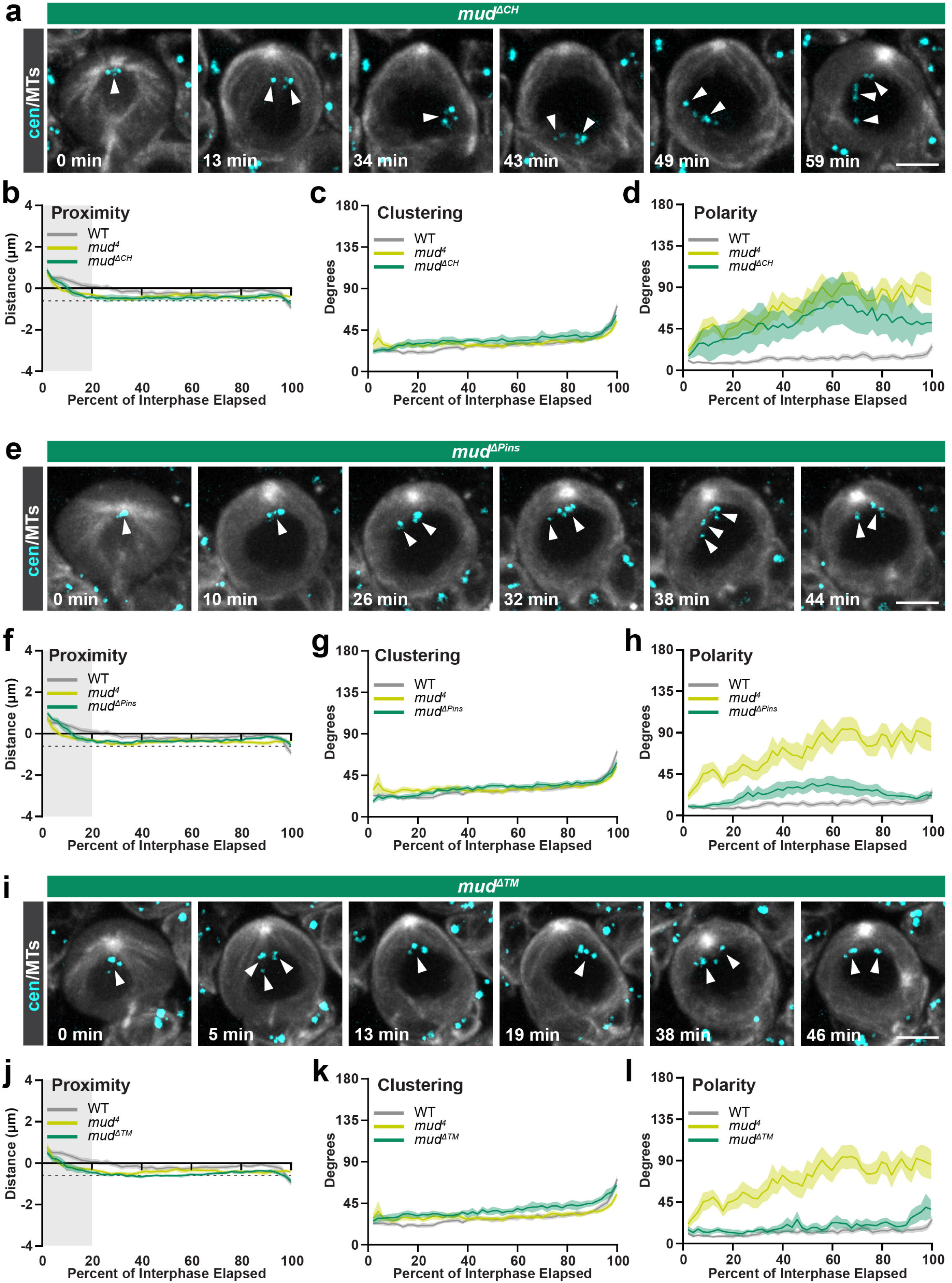
The Mud Calponin homology (CH) domain is required for NB centromere positioning. Image series of representative **(a)** *mud*^Δ*CH*^ (Calponin homology domain), **(e)** *mud*^Δ*Pins*^ (Pins binding domain), and **(i)** *mud*^Δ*TM*^ (Transmembrane domain) mutant third instar larval NB expressing worGal4, UAS-mCherry::α-Tubulin (white) and centromere marker EGFP::Cid (cyan) from the beginning of nuclear envelope reformation (t=0) through prophase. White arrows indicate the positions of centromeres. Graphs of the **(b,f,j)** distance from the nuclear periphery to the centromeres (dotted line = 15% radial distance; gray bar indicates inaccurate nuclear periphery segmentation), **(c,g,k)** angle between each pair of centromeres, and **(d,h,l)** average angle between the apical centrosome and the centromeres. The lines indicate the average at each timepoint and the shaded areas represent the 95% confidence interval. n = 10 NBs, N=3 brain lobes for *mud*^Δ*CH*^; n = 15 NBs, N=3 brain lobes for *mud*^Δ*Pins*^; n = 11 NBs, N=3 brain lobes for *mud*^Δ*TM*^. Scale bar in (a, e, i): 5 µm.

In NBs, Mud interacts with the apical polarity protein Pins to maintain the correct orientation of the mitotic spindle ^66–68^. To determine if the Mud-Pins interaction is also necessary for centromere positioning, we analyzed *mud* mutants lacking the Pins binding domain (*mud*^Δ*Pins*^ ^71^; Fig. S5a). For the majority of interphase, neither peripheralization nor clustering differed significantly from wild type (Fig. 5e-g; Fig. S5c). However, *mud*^Δ*Pins*^ mutant NBs showed a statistically significant difference in centromere polarization compared to wild-type NBs, although the phenotype was not as pronounced as in *mud*^4^ mutants. While centromeres in *mud*^Δ*Pins*^ mutant NBs occupied positions further from the AC than those in wild type, they moved no further than 90° away from the AC (Fig. 5e, h; Fig. S5c). Since there is a small amount of overlap between the Pins domain and the MT domain (Fig.S5a), the intermediate polarity phenotype may be due to a partial disruption of microtubule binding.

Mud has been shown to localize to the nuclear envelope ^70,73,74^. We thus hypothesized that the transmembrane domain might be required for anchoring Mud in the NE and therefore, might be required for centromere positioning. We measured centromere localization in *mud*^Δ*TM*^ ^71^, which is missing the C-terminal predicted transmembrane domain (Fig. S5a). Centromeres retained peripheral localization (within 15% radial position for the majority of interphase) with a small but significant decrease throughout interphase and remained mostly polarized (Fig. 5i, j, l; Fig. S5d). However, there was a subtle yet significant reduction in clustering (8.2° change in average) (Fig. 5i, k; Fig. S5d).

Based on these results, we conclude that Mud’s interaction with Dynein and presumably its MT-binding are required for centromere polarization, while the TM domain contributes to centromere clustering.

### Developmentally regulated gene loci *hb* and *pen* display characteristic polar localization in the NB nucleus

So far, we have shown that the polarized localization of centromeres is regulated by connecting the NBs’ internal polarity axis – orchestrated in interphase via the apically localized MTOC – with the NPC, the Dynein-Mud complex, and Lamin. It is conceivable that this centromere polarization broadly reflects, and possibly even promotes, a stereotypic organization of the NB genome architecture. If so, other chromatin regions might also display stereotypic positioning, such as their proximity to the NE, or their position relative to the NB’s polarity axis. In embryonic neuroblasts, it was shown that the *hunchback* gene (*hb*; Ikaros/Ikzf1 in mammals), encoding for a transcription factor specifying early born neurons ^75^, moves to the neuroblast nuclear periphery, a repressive subnuclear compartment. This led to the model that neuroblasts undergo a developmentally regulated subnuclear genome reorganization, regulating cell fate decisions ^76^. However, in addition to nuclear envelope proximity, it has not been explored whether gene loci also occupy stereotypic positions in relation to nuclear or cellular landmarks. We thus hypothesized that the location of developmentally regulated genes could be influenced by the neuroblast’s intrinsic polarity axis in interphase. To explore this hypothesis, we performed DNA-FISH labeling experiments of two developmentally regulated genes: *hunchback* and *pendulin* (*pen*; KPNA2 in mammals). *pen* expression is increased in larval neuroblasts and early progeny compared to neurons, while *hb* expression is repressed in embryonic and larval NBs ^77,78^. We synthesized DNA probes against *hb* and *pen* and co-stained NBs with anti-γ-Tubulin, anti-Lamin, and anti-Cid. Subsequently, we measured the location of *hb* and *pen*, quantifying their proximity to the NE, and their relative position to the apical MTOC as well as to centromeres (Fig. 6a, c). Consistent with our live cell imaging results (Fig. 1e), the angle between centromeres and the apical centrosome remained below 30 degrees for most wild-type centromeres (Fig. 6a, d). We next measured the angle between the apical centrosome and the gene loci *hb* and *pen* in wild-type NBs. *pen* and *hb* showed more variability in polarization than centromeres. However, both *hb* and *pen* were localized at distinct angular positions relative to the apical centrosome (*hb* mean= 69° SD = +/-25°; *pen* mean= 98° SD = +/-29°) and the average centromere position (*hb* mean= 68° SD = +/-25°; *pen* mean= 97° SD = +/-30°) (Fig 6d-f). *hb* and *pen* also showed more variability in their position relative to the NE. Overall, *hb* and *pen* were significantly further away from the NE compared to centromeres in wild-type larval neuroblasts (Fig. 6a, g). From these data, we conclude that *hb* and *pen* occupy specific spatial regions in the nucleus relative to the cell’s polarity axis and are not as close to the NE as centromeres.

**Figure 6:**
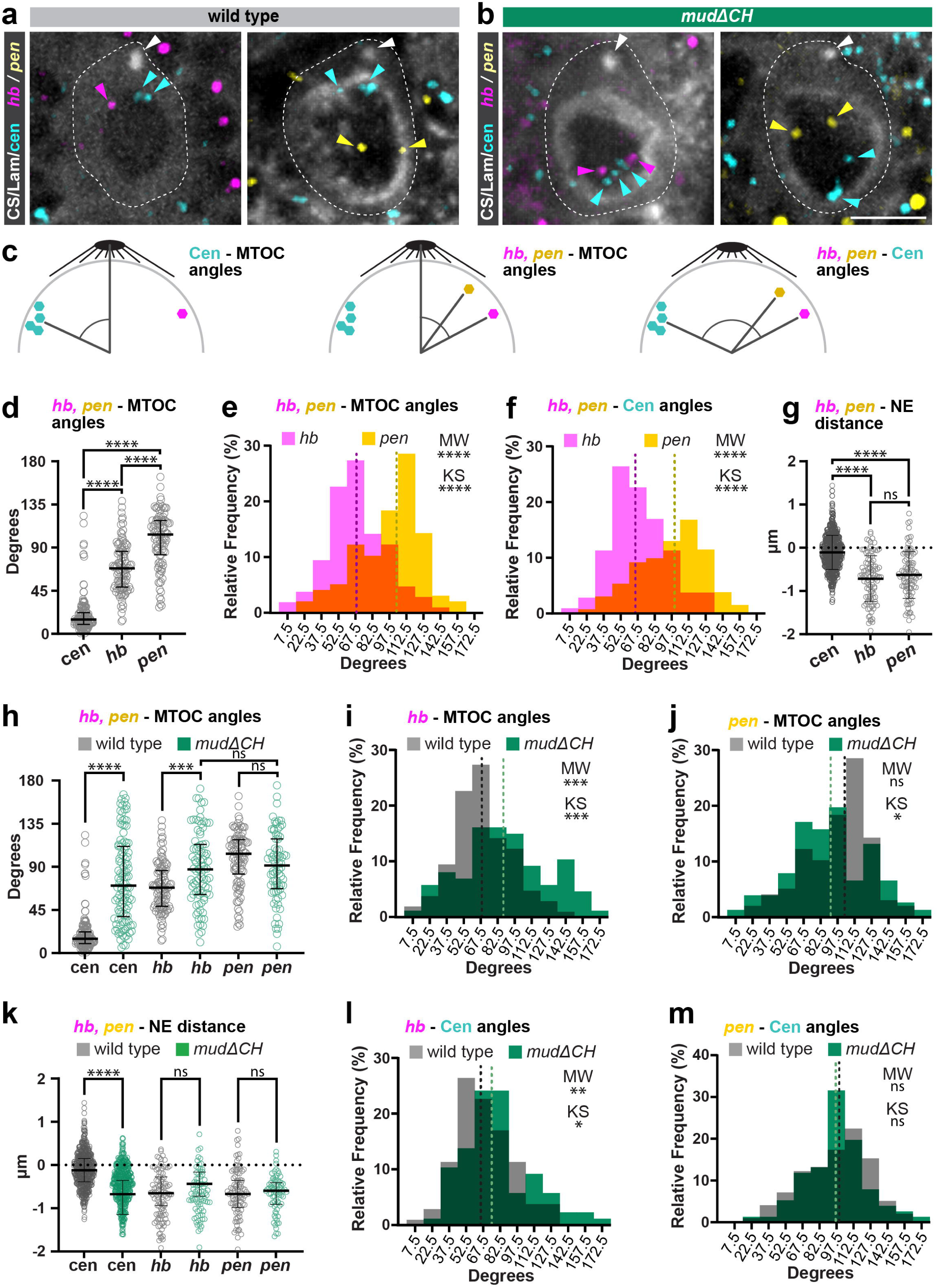
Disruption of centromere tethering impacts *hb* and *pen* polarization. Representative **(a)** wild-type and **(b)** *mud^ΔCH^* third instar NB stained for Lamin (white), the MTOC marker γ-Tubulin (white; white arrowheads), the centromere marker Cid (cyan; cyan arrowheads), and *hb* (magenta; magenta arrowheads) or *pen* (yellow; yellow arrowheads). **(c)** Schematics of measurements used to calculate cen, *hb,* or *pen* positioning. **(d)** Scatterplot of the angles between the apical centrosome (MTOC) and centromeres, *hb*, or *pen* in wild type NBs. Histogram of the angle between *hb* (magenta) or *pen* (yellow) and **(e)** the apical centrosome (MTOC) or **(f)** the centromeres in wild-type NBs. **(g)** Scatterplot of the shortest distance between the nuclear envelope (NE) and centromeres, *hb*, or *pen* in wild type NBs. **(h)** Scatterplot of the angles between the apical centrosome (MTOC) and centromeres, *hb*, or *pen* in wild-type (gray) or *mud*^Δ*CH*^ mutant (green) NBs. Histograms of the angle between the apical centrosome (MTOC) and **(i)** *hb* or **(j)** *pen* in wild-type (gray) or *mud*^Δ*CH*^ (green) mutant NBs. **(k)** Scatterplot of the shortest distance between the nuclear envelope (NE) and centromeres, *hb*, or *pen* in wild-type (gray) or *mud*Δ*CH* mutant (green) NBs. Histogram of the angle between the centromeres and **(l)** *hb* or **(m)** *pen* in wild-type (gray) or *mud*Δ*CH* mutant (green) NBs. Statistical tests used: Mann-Whitney (MW) test in (d-m). Kolmogorov-Smirnov (KS) test in (e, f, i, j, l, m) **** = p<0.0001, *** = p<0.001, ** = p<0.01, *=p<0.05, ns = not significant if p≥0.05. For scatter plots: Center bars denote median and brackets indicate the interquartile range. Dotted line denotes the median in histograms. Scale bar is 5 μm

### Disruption of centromere tethering impacts *hb* and *pen* polarization

To test our hypothesis that centromere positioning might influence the spatial genome organization in the rest of the nucleus, we measured the position of *hb* and *pen* in *mud*^Δ*CH*^ mutants. We used *mud*^Δ*CH*^ because the loss of polarity is highly penetrant throughout interphase (Fig. 5d; Fig. S5b). Furthermore, given that *mud*^Δ*CH*^ mutants only contain a relatively small (153 amino acid) N-terminal deletion, we expect this allele to have fewer potential off-target effects than *mud*^4^ mutants. Compared to wild-type NBs, *hb’s* position in relation to the apical centromere was significantly changed in *mud*^Δ*CH*^ mutants (Fig. 6b, h, i). In contrast, *pen’s* average angle was not significantly altered, although its angular distribution showed a minor difference compared to wild type (Fig. 6b, h, j). This indicates that *pen’s* locus position in relation to the apical MTOC is more varied. Both *hb’s* and *pen’s* position relative to the NE remained unchanged in *mud*^Δ*CH*^ mutants compared to wild-type NBs (Fig. 6k). We conclude that the position of the *hb* locus, and to a lesser degree the *pen* locus, is changed in *mud*^Δ*CH*^ mutants in relation to the neuroblast’s intrinsic polarity axis.

### *hb* positioning in relation to centromeres is also changed in the absence of centromere– MTOC connections

Disconnecting centromeres from the apical MTOC could induce global changes in chromatin organization. Alternatively, centromeres*, hb,* and *pen* could simply change position in relation to the apical MTOC without altering the underlying chromatin architecture. For example, if the nucleus were to rotate after centromeres disconnect from MTOCs while leaving chromatin organization unchanged, centromeres, *hb,* and *pen* would simply change their angular position relative to the apical MTOC without altering their position relative to each other. To test this hypothesis, we measured the angles between centromeres and *hb*, as well as centromeres and *pen* in wild-type and *mud*^Δ*CH*^ mutant NBs. We found that the average angle between centromeres and *hb* was significantly different, whereas *pen* remained unchanged in *mud*^Δ*CH*^ mutant NBs compared to wild type (Fig. 6l, m). We conclude that in *mud*^Δ*CH*^ mutants, the *hb*, but not the *pen* locus, changes its position in relation to NB centromeres. This suggests that disconnecting centromeres from the apical MTOC might change the organization of some chromatin regions.

## Discussion

Spatial genome organization is evolutionary conserved and ensures correct gene expression, necessary for normal development. Disrupting genome architecture can lead to the repositioning of disease-specific genes, potentially changing their activity ^79^. However, how dynamic changes in cell architecture, imposed by cytoskeletal organization and activity, impact chromatin organization is largely unknown. Here, we investigate the mechanism and function of one understudied component of genome organization – centromere positioning – in *Drosophila* third instar larval NBs. We had previously shown that in these neural stem cells, centromeres remain localized near the apical centrosome during interphase and that this localization is dependent on microtubules ^80^.

Here, we provide additional mechanistic insight into the regulation of centromere positioning. Our detailed live-cell imaging analysis pipeline revealed that in interphase neural stem cells, centromeres are located in close proximity to the nuclear envelope, form clusters, and are positioned in a polarized fashion relative to the neuroblast’s apical centrosome. This stereotypic centromere localization is progressively lost in differentiating neuroblast progeny and requires several Nucleoporins (likely as part of the NPC), Dynein, the NuMA-like protein Mushroom body defect, and Lamin.

We propose that on the cytoplasmic side of the NE, microtubules from the apical centrosome are connected to the NPC via Dynein and Mud (Fig. 7). NPCs have been previously shown to interact with Dynein, to be connected to microtubules, and to be linked to chromatin ^81–83^. In several systems, Dynein has been shown to bind to Mud to regulate spindle orientation during mitosis^65^. While Dynein is a minus-end directed microtubule motor protein, Mud interacts with cell-cortex associated proteins, thereby establishing a force-generation machinery that pulls and orients the mitotic spindle ^65^. In mitotic neuroblasts, Mud is concentrated at centrosomes and the apical cortex but is also present at lower levels at the basal cortex ^68^. While we have been unable to detect Mud on the nuclear envelope, Mud has been reported to localize to the interphase nuclear envelope in *Drosophila* S2 cells. Similarly, some Mud splice isoforms have been shown to localize to the nuclear envelope in *Drosophila* male and female gametes ^70,84^. Mud has also been detected at the nuclear envelope in the germarium, including dots on the cytoplasmic side of the nuclear envelope juxtaposed to centromeres ^85^. Thus, Dynein could bind to the NPC via Mud on the outside of the NE, thereby establishing a pulling force as Dynein attempts to walk towards the apical centrosome. Alternatively, Mud could act as a Dynein recruitment, retention, and/or activation factor. Both models are consistent with our finding that Mud’s CH domain, which directly binds to Dynein’s light intermediate chain and is postulated to serve as an activator of cytoplasmic Dynein ^72^, is necessary for polarized centromere localization. Interestingly, the *mud^ΔTM^* domain mutant showed a weak clustering phenotype, without compromising centromere polarization. Although this suggests that the TM domain is not required for centromere polarization, the phenotype is difficult to reconcile with the *mud*^4^ null mutant, which shows no clustering but a polarization phenotype.

**Figure 7:**
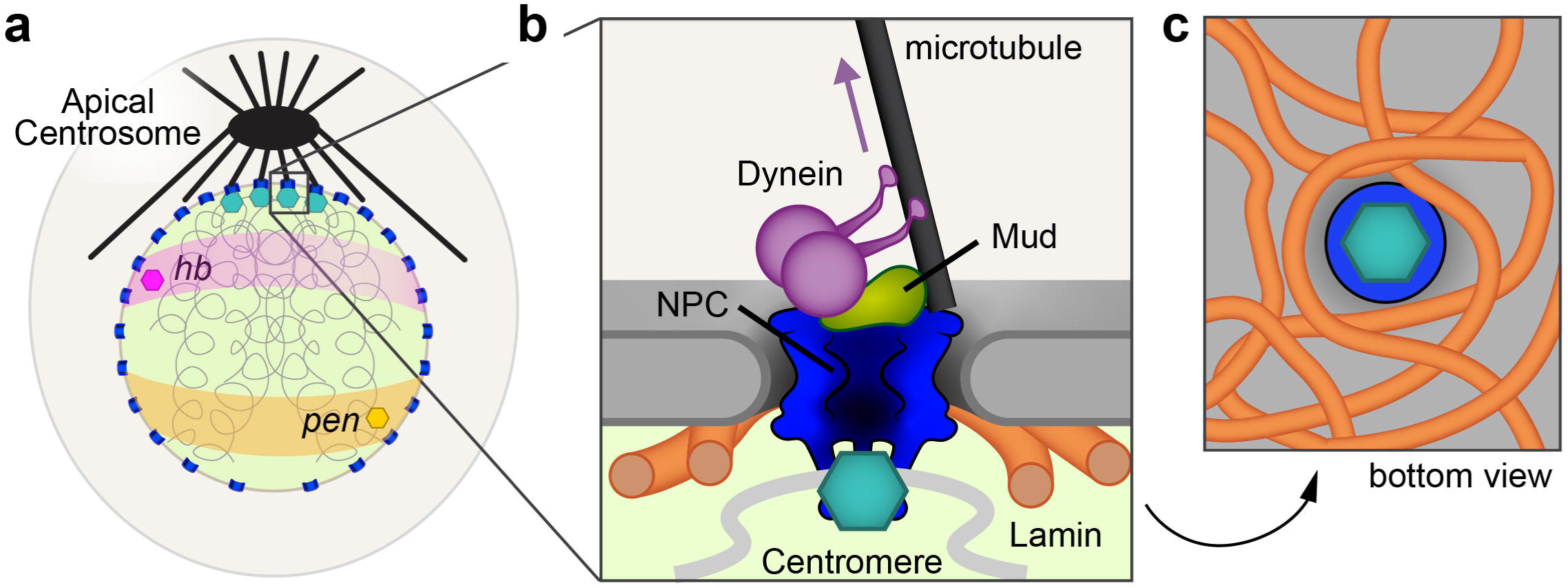
model. **(a**) In third instar larval interphase NBs, microtubules from the apical centrosome (black) connect to both the apical cortex and the apically facing side of the nuclear envelope. Apically enriched Nup62+ NPCs (blue) tether and cluster centromeres (teal) to the apically facing side of the NE. This chromatin organization (chromatin fibers are depicted with curled gray lines inside the nucleus) also impacts the location of developmentally regulated genes such as *hb* (magenta) and *pen* (yellow), which are located in defined positions in relation to the cell’s polarity axis. (**b**) Magnified view corresponding to the boxed region in (a) showing the proposed molecular complex that connects apical MTOC microtubules to centromeres. The purple arrow indicates the direction of movement of Dynein, and chromatin is shown as a curved gray line. **(c)** A view looking from the nucleoplasm up toward the nuclear envelope showing the meshwork of Lamin proposed to anchor NPC positions.

We also observed a modest disruption of centromere polarity in the *mud*^Δ*Pins*^ mutant. The Pins-Mud interaction, mediated by the Mud Pins domain, is necessary for correct spindle orientation in mitotic neuroblasts ^68^. Most likely, this phenotype results from the truncation of the microtubule-binding domain rather than from a loss of interaction with Pins. Further investigations will be necessary to understand the detailed mechanism of Mud’s role in centromere clustering and polarization.

Based on our EM data, it is unlikely that MTs penetrate into the NE to connect with centromeres. Consistent with our data, we propose that on the nucleoplasmic side of the NE, Lamin provides a molecular meshwork, necessary to maintain polarized centromere positioning (Fig. 7).

Non-random centromere positioning has also been observed in other systems, often involving similar molecular players. For instance, Dynein and Mud were both associated with centromere behavior in 8-cell cyst *Drosophila* germline cells. In this system, Dynein is required for centromere grouping and centromere movement. However, in contrast to NBs, in this system, Dynein promotes centromere grouping and movements ^85^. Thus, the finding that Mud is required for non-random and polarized centromere positioning in neuroblasts points to a new function of this universal adaptor protein.

Centromere positioning near an MTOC also occurs in yeast and in the amoeba *Dictyostelium discoideum*. However, in these cases, centromere positioning has been shown to require a SUN-domain protein (Sad1 in yeast, Sun1 in *Dictyostelium*) ^86^. In contrast, we show that in *Drosophila*, SUN- and KASH-domain proteins are not required for polar centromere positioning. Interestingly, both Nups and the Lamin-like proteins CRWN1 and CRWN4 are required for centromere positioning in *Arabidopsis thaliana*. However, in *Arabidopsis*, Nups are required for the scattering rather than for anchoring centromeres at their post-mitotic position ^28,29,87^. Lamin and the Lamin-like CRWN proteins form a lattice-like structure about NPCs that can anchor their positions. Nups have also been shown to directly bind Lamin ^62,63,88,89^.

Surprisingly, none of the identified proteins here dramatically impacted the relative position of centromeres to the nuclear envelope. In all analyzed mutants, centromeres were still localized within 15% of the nuclear periphery, similar to wild-type neuroblasts. This indicates that either centromere proximity is regulated by another mechanism or that several proteins act redundantly to maintain centromeres close to the NE.

Here, we also show that polarized centromere positioning correlates with the stereotypic localization of two developmentally important genes, *hb* and *pen*, which occupy characteristic positions far from centromeres. In *Drosophila* embryonic neuroblasts, *hb* is downregulated after the second NB division, and is subsequently repositioned to the nuclear lamina to enforce gene silencing and prevent erroneous reactivation later ^78^. In contrast, *pen* is expressed in larval NBs and young progeny (INPs and GMCs) but is downregulated in neurons ^77^. Surprisingly, we found that in larval neuroblasts, both *hb* and *pen* occupy stereotypic positions in relation to the neuroblast intrinsic polarity axis and to centromeres. Disconnecting centromeres from the apical MTOC causes *hb* and, to some degree, also *pen* to lose their positional niche relative to the apical centrosome. *hb*, but not *pen*, also changes its position relative to centromeres. This suggests that disconnecting centromeres from the apical centrosome not just induces nuclear rotation but changes chromatin organization, at least to some degree. However, both *hb* and *pen* retained their relative distance to the nuclear envelope, suggesting that other mechanisms are also involved in the peripheral positioning of these loci. These results demonstrate that chromatin organization has far-reaching impacts on spatial genome organization in NBs. Whether and how the polarized positioning of *hb* and *pen* correlates with their gene activity remains to be determined.

Overall, this study demonstrates that intrinsically polarized interphase neuroblasts relay their polarization through their single active, apically localized MTOC across the nuclear envelope to organize chromatin architecture. Our work thus highlights a novel mechanism that translates the cell’s intrinsic polarity axis into genome organization and architecture. In the future, it will be interesting to see whether polarity-dependent chromatin organization also occurs in other systems and how it impacts cellular functions under physiological conditions.

## Methods

### Fly strains

Mutant alleles, transgenes, and fluorescent markers:

*worGal4, UAS-mCherry::*α*-Tubulin/CyO* ^35^, *Cid-mRFP1 II.1, Cid-mRFP1 II.2* ^90^, *Cid-EGFP III.2* ^37^, *GFP-cid* (BDSC:25047), *UAS-Cherry::Jupiter* ^91^, *koi*^80^ (BDSC:25105), *koi* deficiency *Df(2R)Exel6050* (BDSC:7532), *koi* RNAi line *P{TRiP.HMS02172}attP40* (BDSC:40924), *klar^Marb-CD4^* ^92^ (BDSC:25097), *klar* deficiency *Df(3L)emc-E12/TM6B* (BDSC:2577), *klar* RNAi lines *P{TRiP.JF02944}attP2 e1*, P{TRiP.HMS01612}attP2, *klar^GD9271^*, (BDSC: 28313, 36721, VDRC:32836), *Msp300*^Δ*KASH*^ ^56^, *Msp300* deficiency Df(2L)Exel6011 (BDSC:7497), *cnb^GD11735^* RNAi line (v28651) ^93^, *Nup54* RNAi line *Nup54^KK102105^*(VDRC:103724), *Nup98-96* RNAi line *Nup98-96^GD6897^* (VDRC:31198), *Nup62* RNAi line *P{TRiP.GLV21060}attP2* (BDSC:35695), *P{EGFP.Nup62}III.1* (BL95381), *Nup62^ex161^/CyO* (BDSC:95380), *w[*]; P{w[+mW.hs]=FRT(w[hs])}G13* (BL1956), *UAS-Lamin::GFP* (BDSC: 7377, 7378) ^40^, *Lam^A25^* (BDSC: 25092) ^94^, *Lam^04643^* (BDSC:11384) ^95^, *Dhc64C* RNAi line *Dhc64C^GD12258^*(VDRC:28054), *mud^4^* (BDSC:9563) ^96^, *mud*^Δ*CH*^*, mud*^Δ*PINS.GFP*^ *, mud*^Δ*TM.GFP*^ ^71^.

### Mosaic analysis with a repressible cell marker (MARCM)

For MARCM clones, we recombined *nup62^ex161^* onto *FRTG13* (BDSC:1956), and crossed either BDSC:1956 or the recombinant to the following line: *yw, hsFLP70; tubP-Gal80, FRTG13/(CyO); worGal4, UAS-mCherry::*α*-Tubulin/TM6B.* Seperately, we also recombined *cid::mRFP* onto *FRTG13* and crossed with *yw, hsFLP70; tubP-Gal80, FRTG13/(CyO); worGal4, UAS-Lamin::GFP/TM6B*. Recombination was induced by incubation at 37°C for 1 hr.

### Immunohistochemistry

The following antibodies were used: mouse anti-γ-tubulin (1:2000, Millipore Sigma, catalog number SAB4701044), mouse anti-Lamin Dm0 (1:50, Developmental Studies Hybridoma Bank, ADL67.10), rat anti-Cid monoclonal 4F8 (1:400, Fisher Scientific catalog number 50-199-3653), guinea pig anti-Lamin Dm0 (1:5000, kind gift from Veena Parnaik), guinea pig anti-Asterless (1:20,000, kind gift from J. Raff, 1:20,000), rabbit anti-centrosomin (1:1000; kind gift from Tim Megraw).

Larval brains (72 to 96 h after egg laying) were dissected in Schneider’s insect medium (Sigma, catalogue no. S0146-100ML) for no more than 30 min (see ^97^ for detailed dissection instructions). Dissected samples were fixed in 4% paraformaldehyde in Schneider’s medium for 20 min on a rotator at room temperature. After fixing, the samples were washed at least three times with PBSBT (1X PBS, 0.1% vol/vol of Triton-X-100, and 1% wt/vol BSA) for 20 min each. Primary antibody dilution was prepared in 1X PBSBT, and samples were incubated in primary antibody solution for at least 1 d at 4°C with agitation. After primary labeling, samples were washed with PBSBT at least three times for 20 min each. Secondary antibody solution was prepared in 1X PBSBT, and samples were covered and again incubated for at least 1 d at 4°C with agitation. After incubation, the samples were washed three times with PBST (1X PBS, 0.1% vol/vol of Triton-X-100). Mounting slides were prepared ahead of time by affixing two glass coverslips onto a glass slide with nail polish to form a thin channel to hold the sample. Final dissections to remove the brains from the cuticles were done in PBSBT, and the brains were then transferred via pipette to equilibrate in ProLong Diamond mounting media (ThermoFisher Scientific, catalogue number P36961). After equilibration, samples were transferred to the previously prepared slides and sealed with a glass coverslip and nail polish.

### Live-cell imaging

72 to 96 h after egg laying old larval brains were dissected using microdissection scissors (Fine Science Tools, catalogue no. 15003-08) and forceps (Dumont #5, Electron Microscopy Sciences, item number 0103-5-PO) in Schneider’s medium supplemented with 1% bovine growth serum (HyClone, item number SH30541.03) and transferred to chambered slides (Ibidi, catalogue no. 80826) for imaging. Detailed dissection and live cell imaging sample preparation instructions can be found here ^97,98^.

Both live and fixed samples were imaged with an Intelligent Imaging Innovations (3i) spinning disk confocal system, consisting of a Yokogawa CSU-W1 spinning disk unit and two Prime 95B Scientific CMOS cameras and a 60x/1.4NA oil immersion objective. Voxels are 0.22 × 0.22 × 1 µm. Temporal resolution was 30 s per frame.

### Transmission Electron Microscopy

Larval brains were fixed in ½ strength Karnovsky’s fixative (2.5% glutaraldehyde, 2% paraformaldehyde in 0.1M sodium cacodylate buffer, pH 7.3) overnight at 4 °C. Fixed samples were rinsed with 0.1M cacodylate buffer, post fixed with 1% osmium tetroxide/0.8% potassium ferricyanide for 1 h at 4 °C, rinsed with cacodylate buffer, and incubated in 0.2% tannic acid for 15 min, rinsed with water and en bloc stained with 1% uranyl acetate for 1 h. The samples were then rinsed with distilled water and dehydrated through a graded series of alcohols followed by propylene oxide and embedded in Eponate12 resin (Ted Pella, Inc, Redding, CA). 70nm ultrathin sections were cut using a Leica EM UC7 ultramicrotome, contrasted with uranyl acetate and lead citrate, and imaged on a ThermoFisher Talos L120c transmission electron microscope at 120kV. Digital images were acquired with a Ceta 16M CMOS 4kx4k digital camera system.

### Photoactivatable Microtubule Inhibitor

Brains were treated with 30uM SBTub2M ^53^, 1% DMSO final concentration for 30 minutes prior to imaging. SBTub2M was activated by imaging through the z-stack with 405 nm 40% laser exposure for 0.5 s per plane.

### DNA-FISH

DNA-FISH probes were generated using the Invitrogen FISH Tag Alexa Fluor 488 DNA Kit (Catalogue #F32947) according to manufacturer instructions. The following primers were used to amplify DNA for the probes:

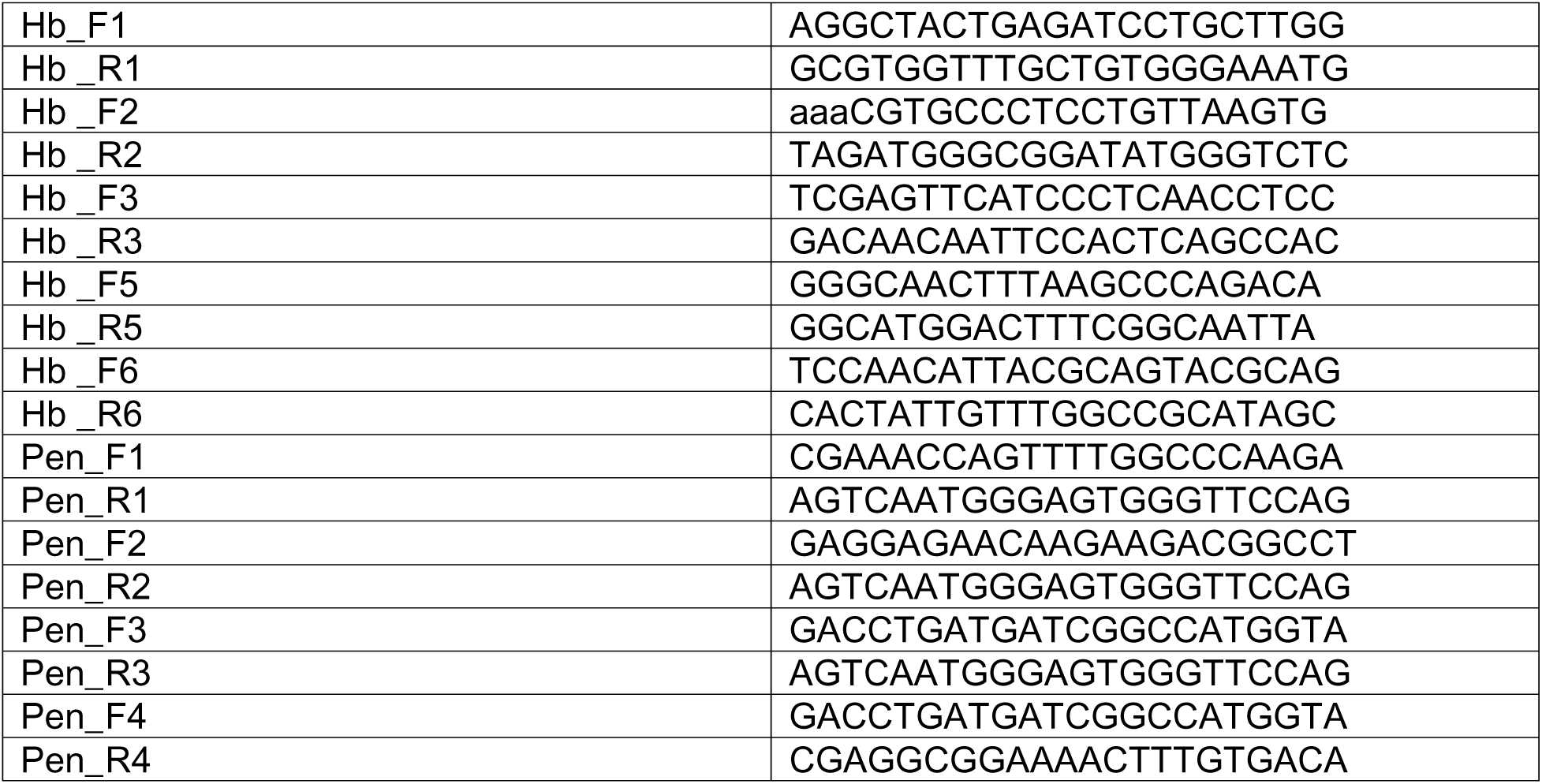

DNA-FISH was performed based on a published protocol ^99^ with small modifications. In brief, brains were dissected and fixed as for immunofluorescence (above). They were then washed 3 times in PBS + 0.1% Tween 20 for 20 m each, permeabilized 1 hr in PBS + 0.3% Triton X-100, and transferred through a graded series of 15 m washes into 50% formamide, 4X saline-sodium citrate (SSC) buffer, 100 mM NaH_2_PO_4_, pH 7.0, 0.1% Tween 20, incubated for 30 min at 37°C, then denatured for 15 min at 80°C. 80-100 ng probe in 50% formamide, 10% dextransulfate, 2X SSC, 0.5 mg/ml salmon sperm DNA was denatured 10 min at 90°C. As much supernatant as possible was removed from brains and probe was added, flicked to mix, and incubated at 91°C for 2 min. Hybridization was performed overnight at 40 °C in a thermomixer with gentle agitation (500 rpm). Brains were then transferred back through a graded series of 15 min washes into PBS + 0.1% Tween 20 and post-fixed with 4% paraformaldehyde for 30 min. Immunofluorescence staining and imaging was performed as described above.

### Analysis

All analyses were done in Imaris 10.2 and ImageJ 1.54. Imaris was used to segment nuclei and centrosomes by training pixel classifiers.

### Neuroblast nuclei segmentation and measurements

To segment the NB nuclei of brains expressing mCherry::α-Tubulin, we trained a pixel classifier on brains expressing both mCherry::α-Tubulin and UAS, Lamin::GFP. The pixel classifier was only trained on the Tubulin signal; the Lamin signal was used visually by the authors to ensure correct training pixels were selected. The nuclei used were manually edited to remove any protruding artefacts from the surface. For segmenting neuroblast centrosomes, a baseline pixel classifier was generated using cells expressing mCherry::α-Tubulin and was updated for individual cells to ensure accurate segmentation of the centrosome. Centromere (Cid) spots were segmented using the Spots tool with a size of 0.4 µm, quality filtered, and spots corresponding to noise or hot pixels were removed manually.

For centromere positioning analysis in NBs, we selected nuclei that were well segmented for a complete interphase. Only one division was analyzed per cell. To normalize differing interphase lengths, we calculated the percentage of interphase elapsed (T_x_/(T_end_-T_beginning_), where T_x_ is a given timepoint; T_end_ is the time of nuclear envelope breakdown, defined as the last frame when the nucleus was able to be accurately segmented; T _beginning_ is the first frame of interphase, defined as when the nucleus was first able to be segmented). We binned time measurements into buckets at 2% intervals, with 50 total buckets.

We used the built in Shortest Distance to Surfaces measurement in Imaris to measure the shortest distance from each centromere spot to the nuclear periphery (peripheralization). To calculate the average angle among centromeres at a given timepoint (clustering), we measured the absolute value of the angles between each unique pair of centromeres and then averaged those angles. We used the positional coordinates of the centroid of the nucleus, calculated by Imaris, as the vertex of these angles and the positional coordinates of the centromere spots as the terminal points of the angle. We then calculated the vectors from the vertex to each terminal point. This and all other angles below were calculated from vectors *a*, *b* using the following formula: cos^-1^(*a*•*b*/|*a*||*b*|). Angles calculated from this formula take on absolute values between 0° and 180°, inclusive. To calculate the average angle between the centromeres and the apical centrosome (polarization), we used the average of all centromere positions as one terminal point, the centroid of the apical centrosome as the other terminal point, and the centroid of the nucleus as the vertex.

### Clonal nucleus segmentation and analysis

To segment the NB and progeny nuclei of brains expressing UAS, Lamin::GFP, we trained a pixel classifier using the Lamin signal. We were unsuccessful training a pixel classifier that included the lamin signal at the periphery of the nucleus, so segmentations sit inside the lamin signal. This results in a segmentation that underestimates the position of the nuclear periphery. To estimate the approximate decrease in nuclear radius, we measured the distance from the edge of the segmented nucleus to the center of the Lamin signal.

Peripheralization and clustering measurements were calculated as for NB nuclei in the section above. For polarization, we measured the change in position over the course of 3 minutes as the angle between the vector from the T_0_ nucleus centroid position to the T_0_ position of a given centromere and the vector from the T_3_ nucleus centroid position to the T_3_ position of that same centromere. We used spots tracking in Imaris to track individual centromere spots over time.

### EGFP::Nup62 localization analysis

For quantification, nuclei were segmented with a pixel classifier using EGFP::Nup62 signal in Imaris. Spots were generated using the EGFP::Nup62 signal and the segmented nuclei were used to select 0.4 µm spots localized at the nuclear periphery and separately spots within the nucleus. We also generated spots at centromere foci as above. We selected NBs that completed one interphase and which had centromeres present at approximately the middle z-slice of the cell. We used the 5 middle z-slices of the cell and selected spots on the periphery 0 to +/-15° (apical), 75°-105° and -75° to -105° (lateral) and +/-165° to 180° (basal) from the average centromere position (using the nucleus centroid as the vertex). At each timepoint, the average intensity of spots (background signal) inside the nucleus was subtracted from the selected spots. Spots for each location were used if there were at least 3 spots at a given timepoint. These spots were averaged at each timepoint and ratios were calculated. To test for significance, a ratio t-test was used to test the null hypothesis that each ratio = 1.

### DNA-FISH Analysis

Spots were generated using the Spots tool in Imaris for γ-Tubulin, Cid, *hb*, and *pen* signals and were quality filtered. The brightest 1-2 spots per cell were selected for *hb* and *pen* (the selected spots were notably brighter than any other spots). Background signal in the cytoplasm but not nucleoplasm plus nuclear peripheral signal in the 405 channel stained with anti-γ-Tubulin and anti-Lamin Dm0 allowed us to train a pixel classifier to segment the nuclei. The nuclei used were manually edited to remove any protruding artefacts from the surface. We used the average position of Cid spots within a cell and individual positions of *hb* and *pen* loci for the angle terminal points and the centroid of the nucleus as the vertex. Angles were calculated as described above. The measurement for the distance between the gene loci and the nuclear periphery was calculated in Imaris.

### Graphing, Statistics, and Display

Graphing and statistics were performed using GraphPad Prism 10 and figures were constructed using Adobe Photoshop 2025 and Adobe Illustrator 2025. Supplemental movies were exported from Imaris and annotated using Adobe Premiere 2025.

## Acknowledgements

We thank Tim Megraw for the anti-Cnn antibody, Veena Parnaik for the anti-Lamin Dm0 antibody, and Jordan Raff for the anti-Asl antibody. The anti-Lamin antibody ADL67.10 developed by Fisher, P. A. was obtained from the Developmental Studies Hybridoma Bank, created by the NICHD of the NIH and maintained at The University of Iowa, Department of Biology, Iowa City, IA 52242. Electron microscopy data were generated using the Fred Hutchinson Cancer Center Electron Microscopy shared resource, which is supported in part by the Cancer Center Support Grant P30 CA015704-40. We further thank Yohanns Bellaiche for the *mud* deletion alleles, Steve MacFarlane for TEM sample processing and Ryan Wilkerson for creating PowerShell scripts for data processing. We also thank members of the Cabernard laboratory for helpful discussions and comments. Fly stocks were obtained from the Vienna Drosophila Resource Center (VDRC, www.vdrc.at) and the Bloomington Drosophila Stock Center (BDSC), which is supported by grant P40OD018537 from the NIH Office of Research Infrastructure Programs (ORIP) in collaboration with the National Institute of General Medical Sciences (NIGMS), National Institute of Neurological Disorders and Stroke (NINDA) and National Institute of Child Health and Human Development (NICHD). This work was supported by the American Cancer Society (PF-22-089-01-DMC to J.A.T) and the National Institutes of Health (R35GM148160 to C.C).

---

**Supplemental Figure 1: Centromeres are polarized, clustered, and peripherally localized in interphase NBs.**

Scatterplots showing the **(a)** distance from the nuclear periphery to centromeres. The dotted line represents 15% radial distance. The gray bar indicates inaccurate nuclear periphery segmentation due to low α-Tubulin signal contrast. Negative and positive values indicates that the centromeres are positioned inside or outside the segmented nucleus, respectively. **(b)** Scatterplot showing the average of the angles between each pair of centromeres, and **(c)** angles between the apical centrosome and the average position of the centromeres. The plots show measurements from 18 neuroblasts. Histograms of **(d)** NB nuclear and **(e)** progeny cell interphase radii (n=10 Nbs, 18 progeny cells). The mean is shown with the dotted line. **(f)** Scatterplot of the shortest distance between the nuclear envelope (NE) and centromeres in NBs (gray), young (#1-10, orange), or older progeny (#13-29, purple). Measurements are derived from the representative individual NB clone in Figure 1 over the course of the 3 hr movie. Progeny #11 and 12 divided during the course of the movie and were thus excluded. **(g)** Box plots of the average angle between centromeres (left) or angular speed (Δ Degrees/3 m) (right) for the NB (gray), young (#1-10, orange), or older progeny (#13-29, blue) in the representative individual NB clone shown in Figure 1. **(h, i, j)** Representative images of wild type third instar NBs and progeny cells, expressing worGal4, UAS-mCherry::α-Tubulin and stained for Lamin (purple), Cid (teal), and the MTOC marker Centrosomin, cnn (yellow). Asymmetric MTOCs can be seen in progeny (h,i; top and high magnification images in middle row) adjacent or close to NBs, but not in (j) older progeny. White arrowheads highlight Cnn^+^ MTOCs, teal arrowheads indicate centromeres. White dotted lines indicate the cell periphery. The white bracket (i) indicates a row of progeny between the NB and indicated progeny. Statistical tests used: Mann-Whitney test (f, g) **** = p<0.0001, * = p<0.05, ns = not significant, p≥0.05. Scale bar = 5 µm in (h, i, j); 2 µm in high magnification images in (h, i).

**Supplemental Figure 2: The LINC complex is not required for centromere positioning** Image series of representative **(a)** *koi*^80^ null, **(b)** *koi* RNAi expressing, **(c)** *klar^marb-CD4^* null, **(d, e, f)** *klar* RNAi expressing and **(g)** *msp300*^Δ*KASH*^ mutant third instar larval NBs. In all conditions, the representative NBs also express worGal4, UAS-mCherry::α-Tubulin (white) and the centromere marker EGFP::Cid (cyan). White arrows indicate the positions of centromeres. 0 minutes is defined as the beginning of nuclear envelope reformation. Scale bars = 5 µm.

**Supplemental Figure 3: Centromeres lose polarity in *nup54* RNAi, *nup98-96* RNAi, and nup62^ex161^ null NBs**

Image series of representative **(a)** *nup54* or **(b)** *nup98-96* RNAi expressing third instar NBs. Representative image sequences of **(c)** a wild-type control or **(d)** *nup62^ex161^* homozygous mutant NB clone, labelled with worGal4, UAS-mCherry::α-Tubulin (white) and centromere marker EGFP::Cid (cyan) from the beginning of nuclear envelope reformation (t=0). White arrows indicate the positions of centromeres. Scale bars = 5 µm.

**Supplemental Figure 4: Centromere localization depends on microtubules, Nup62, Lamin, cytoplasmic Dynein, and Mud**

Scatterplots of plotting the distances from the nuclear periphery to the centromeres (left; dotted line = 15% radial distance; gray bars indicate inaccurate nuclear periphery segmentation), angles between each pair of centromeres (middle), and angles between the apical centrosome and the centromeres (right) for **(a)** *cnb* RNAi (dark red, n = 13 cells from N=2 brain lobes), **(b)** *nup62* RNAi (blue, n=13 cells from N=4 brain lobes), **(c)** *Lam^A25^* / *Lam^04643^* mutant (orange; n = 7 cells, N=2 brain lobes), **(d)** *dhc64C* RNAi (purple; n=8 cells, N=4 brain lobes), and **(e)** *mud^4^*null mutant NBs (lime green; n = 12 cells, N=3 brain lobes). Wild-type measurements are shown in gray (n = 18 cells from N=3 brain lobes). In (left), a negative value indicates that the centromere is positioned inside the segmented nucleus, while a positive value indicates that the centromere is outside.

**Supplemental Figure 5: The Mud Calponin homology (CH) domain is required for NB centromere positioning**

Cartoon of the full-length Mud protein (top) with mapped domains. Deletion mutants are shown below. Scatterplots of the distance from the nuclear periphery to the centromeres (left; dotted line indicates 15% radial distance; gray bar highlights inaccurate nuclear periphery segmentation). A negative value indicates that the centromere is positioned inside the segmented nucleus while a positive value indicates that the centromere is outside. Scatterplot showing the average angle between each pair of centromeres for a given timepoint (middle) and angles between the apical centrosome and the average position of the centromeres (right) for b) *mud*^Δ*CH*^ (Calponin homology domain); n = 10 NBs, from 3 different brain lobes, **(c)** *mud*^Δ*Pins*^ **(a)** (Pins binding domain), n = 15 NBs, from 3 different brain lobes, and **(d)** *mud*^Δ*TM*^ (Transmembrane domain) mutant third instar larval NBs; n = 11 NBs, from 3 different brain lobes. Wild type measurements are shown in gray; n = 18 NBs from 3 different brain lobes.

## Movie legends

**Movie 01: wild-type neuroblast expressing worGal4, UAS-mCherry::αTubulin and Cid::EGFP**

The first loop of the movie shows a wild-type third instar larval neuroblast expressing mCherry::αTubulin (white, microtubules) and Cid::EGFP (teal, centromeres). White and teal arrowheads denote the apical centrosome and centromeres, respectively. Note that the centromeres remain near the apical centrosome throughout interphase. The second loop of the movie shows the same interphase overlayed with spots for the centromeres (yellow) and surface reconstructions for the apical centrosome (magenta) and nucleus (purple) that were used for measurements of centromere positioning. Time scale is min:sec, and the scale bar is 2 μm.

**Movie 02: *cnb* RNAi neuroblast expressing worGal4, UAS-mCherry::αTubulin and Cid::EGFP**

The movie shows a *cnb* RNAi third instar larval neuroblast expressing mCherry::αTubulin (white, microtubules) and Cid::EGFP (teal, centromeres). White and teal arrowheads denote the apical centrosome and centromeres, respectively. Note that during early interphase, centromeres remain localized next to the apical centrosome while it is present (46:30). The apical centrosome disappears during interphase and centromeres move away from their previous position (62:30). The apical centrosome reappears as the cell prepares to enter mitosis (83:00). Time scale is min:sec, and the scale bar is 2 μm.

**Movie 03: *nup62* RNAi neuroblast expressing worGal4, UAS-mCherry::αTubulin and Cid::EGFP**

The movie shows a *nup62* RNAi third instar larval neuroblast expressing mCherry::αTubulin (white, microtubules) and Cid::EGFP (teal, centromeres). White and teal arrowheads denote the apical centrosome and centromeres, respectively. Note that during interphase, centromeres move away from the apical centrosome even though it is still present (79:00, 93:30). Time scale is min:sec, and the scale bar is 2 μm.

**Movie 04: wild-type neuroblast expressing EGFP::Nup62 and Cid::mRFP**

The movie shows a wild-type third instar larval neuroblast expressing EGFP::Nup62 (white) and Cid::mRFP (teal, centromeres). White and teal arrowheads denote Nup62 and centromeres, respectively. Note that at the beginning of nuclear envelope reformation, Nup62 preferentially accumulates first at the apical side of the nuclear envelope next to centromeres and subsequently spreads downward (45:00 and 105:30-107:00). Time scale is min:sec, and the scale bar is 2 μm.

**Movie 05: *lam^A25^/lam^04643^*mutant neuroblast expressing worGal4, UAS-mCherry::αTubulin and Cid::EGFP**

The movie shows a *lam^A25^/lam^04643^* mutant third instar larval neuroblast expressing mCherry::αTubulin (white, microtubules) and Cid::EGFP (teal, centromeres). White and teal arrowheads denote the apical centrosome and centromeres, respectively. Note that during interphase, centromeres move away from the apical centrosome even though it is still present (45:00, 66:30). Time scale is min:sec, and the scale bar is 2 μm.

**Movie 06: *dhc64C* RNAi neuroblast expressing worGal4, UAS-mCherry::αTubulin and Cid::EGFP**

The movie shows a *dhc64C* RNAi third instar larval neuroblast expressing mCherry::αTubulin (white, microtubules) and Cid::EGFP (teal, centromeres). White and teal arrowheads denote the apical centrosome and centromeres, respectively. Note that during interphase, centromeres move away from the apical centrosome even though it is still present (92:30). Time scale is min:sec, and the scale bar is 2 μm.

**Movie 07: *mud^4^* mutant neuroblast expressing worGal4, UAS-mCherry::αTubulin and Cid::EGFP**

The movie shows a *mud^4^* mutant mutant third instar larval neuroblast expressing mCherry::αTubulin (white, microtubules) and Cid::EGFP (teal, centromeres). White and teal arrowheads denote the apical centrosome and centromeres, respectively. Note that during interphase, centromeres move away from the apical centrosome even though it is still present (26:00, 40:00). Time scale is min:sec, and the scale bar is 2 μm.

