## Supplementary figures and images for "Asymmetric centromere and gene locus positioning in *Drosophila* neural stem cells"

### Supplemental Figure 1

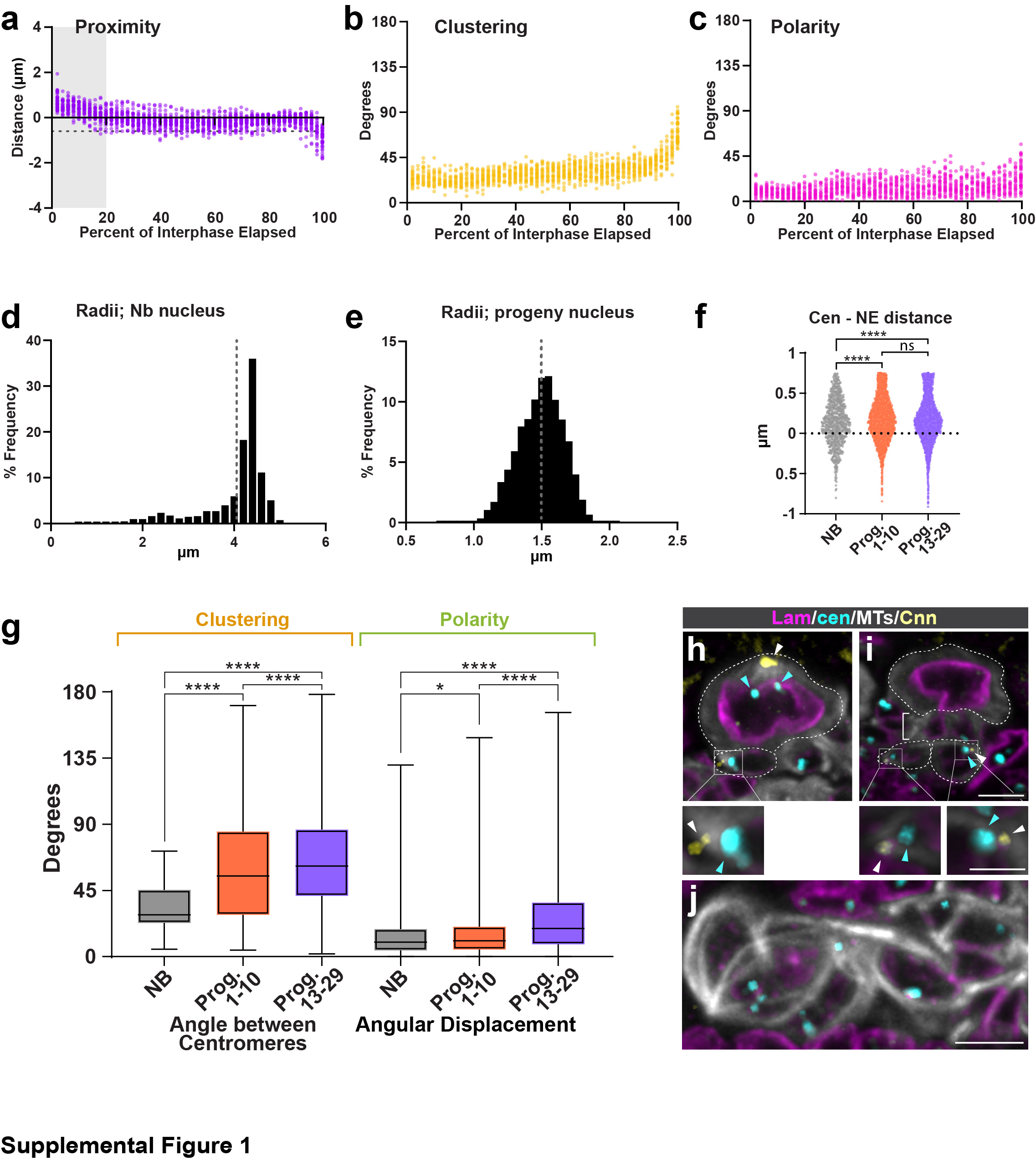

### Supplemental Figure 2

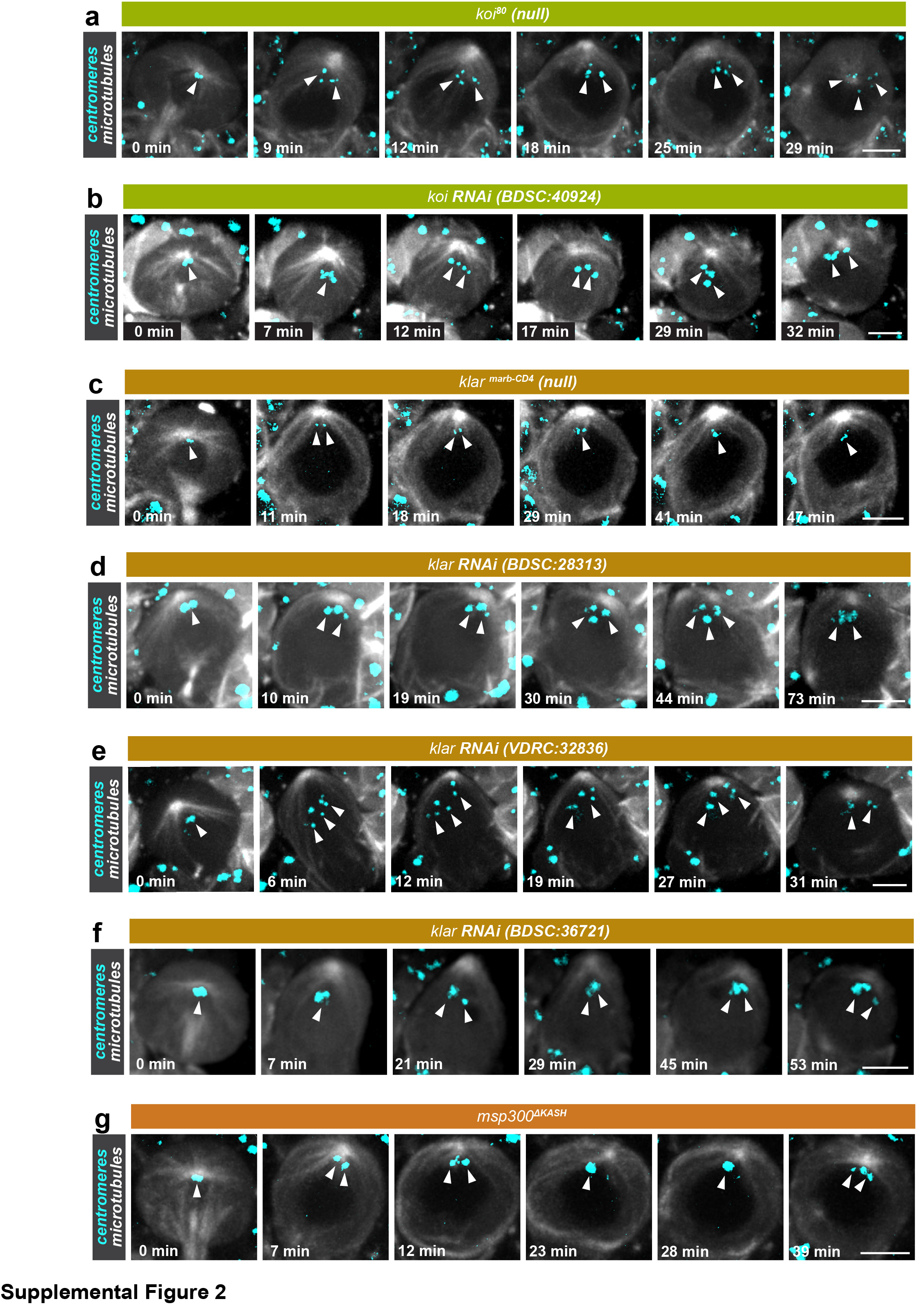

### Supplemental Figure 3

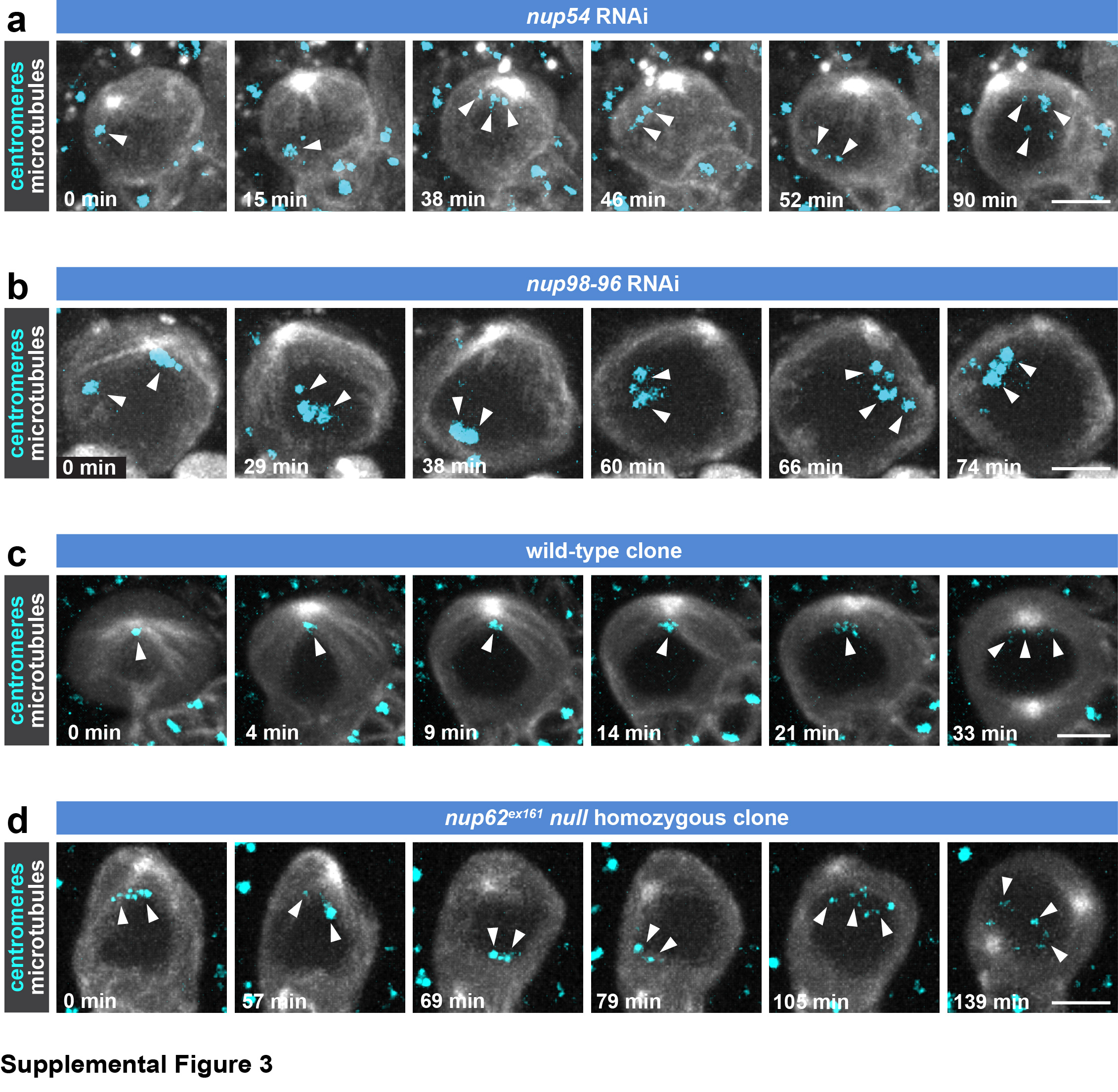

### Supplemental Figure 4

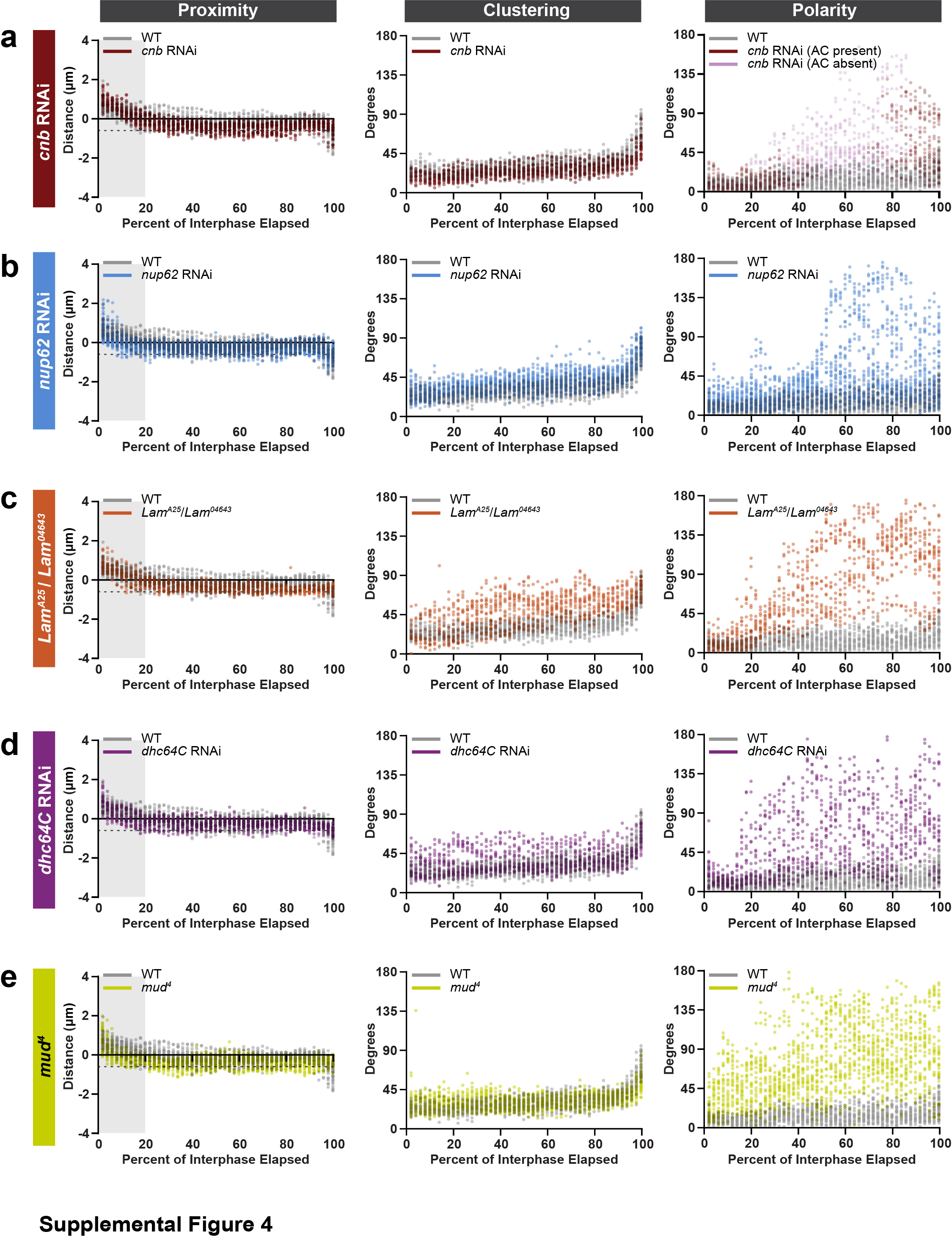

### Supplemental Figure 5

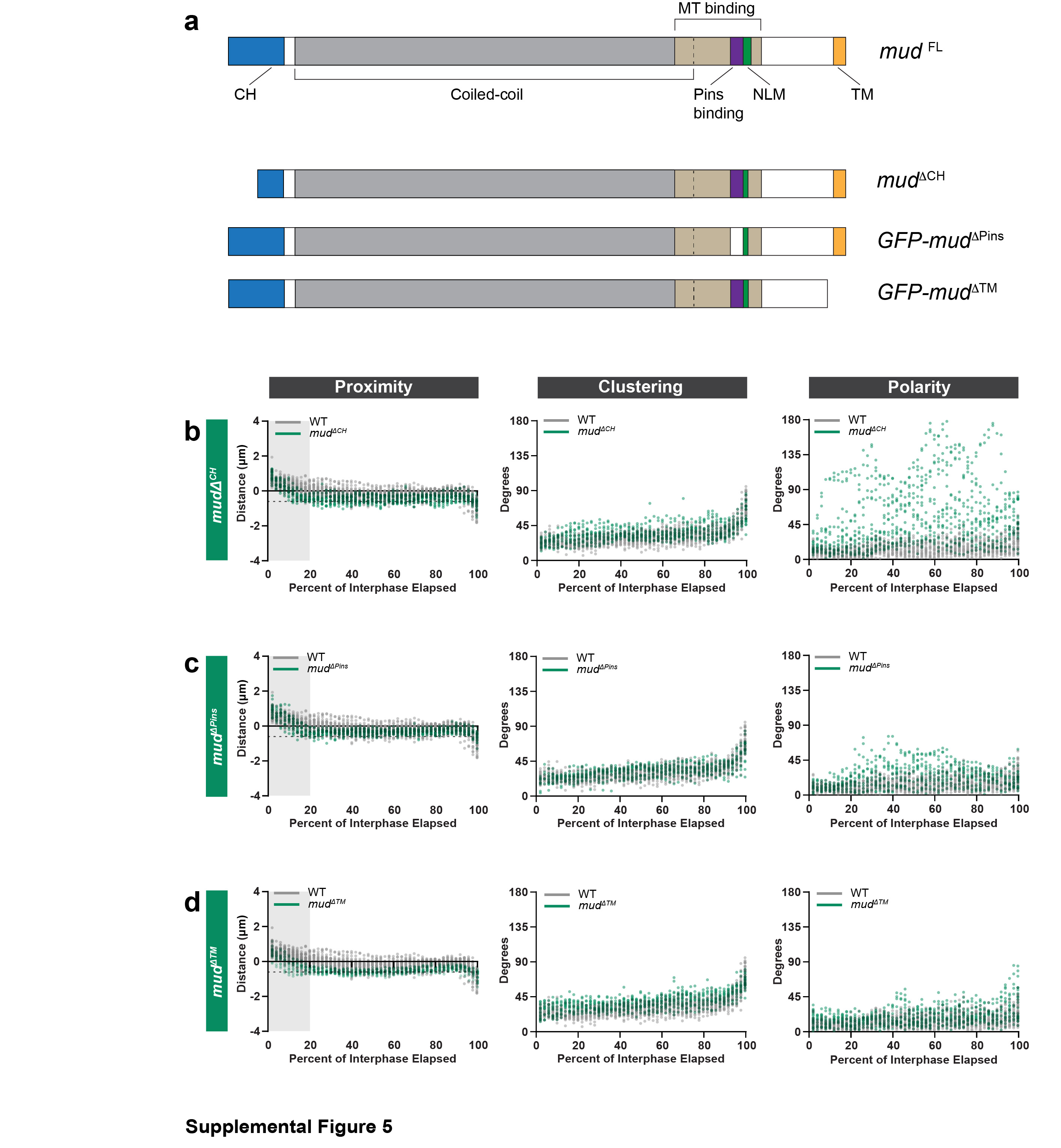
